# Genome Organizer SATB1 selectively activates a defined subset of EMT genes driving metastatic breast cancer

**DOI:** 10.64898/2026.05.29.728584

**Authors:** Yoshinori Kohwi, Yelena Vayn, Mari Grange, Bao Ho, Diane Heditsian, Terumi Kohwi-Shigematsu

## Abstract

SATB1 reshapes chromatin architecture and transcriptional programs to promote breast cancer metastasis. However, its key downstream effectors remain incompletely defined. Here, we aimed to identify actionable drivers of invasion by focusing on epithelial-mesenchymal transition (EMT) genes. We identified 98 of 300 curated EMT-promoting genes as direct SATB1 targets in human breast epithelial cells (MCF10A-1) rendered tumorigenic with metastatic traits by SATB1 transduction, using Global Run-On Sequencing (GRO-seq) to measure nascent transcripts. These SATB1-activated EMT genes regulate extracellular matrix remodeling, hypoxia-responsive transcriptional programs, and tumor microenvironmental programs linking angiogenesis and immune evasion, collectively enhancing metastatic competence. Triple-negative breast cancer (TNBC) is a heterogeneous disease characterized by frequent metastasis and chemoresistance. Among the four TNBC molecular subtypes, the 98 SATB1-regulated EMT genes were significantly enriched and activated in the Basal-like 2 (BL2) subtype (Fisher’s exact test: p = 4.53e-9), which is associated with aggressive behavior, poorer clinical outcomes, and reduced treatment responsiveness. In contrast, SATB1-independent EMT genes showed no enrichment in BL2, indicating selective regulation of EMT genes by SATB1. We further analyzed nascent transcripts induced by the environmental carcinogen benzo[a]pyrene (B[a]P), a known breast carcinogen. Half of the 72 EMT genes activated after short-term B[a]P exposure overlapped with SATB1-dependent EMT genes, indicating that two distinct etiologies, SATB1 and B[a]P, converge on a largely shared network of invasion-promoting genes. These results show that EMT genes are not globally or randomly activated in breast cancer but are selectively activated, defining an EMT gene network associated with metastatic risk. This gene signature may serve as a prognostic marker pending further validation.

## Introduction

Identifying the key drivers of cancer metastasis and elucidating their mechanisms remain major challenges in cancer research. The nuclear architectural protein SATB1 has emerged as a critical regulator of metastasis in epithelial cancers. Initially identified in breast cancer^1–6^, SATB1 was subsequently implicated in multiple other malignancies^7–9^. SATB1 functions as a genome-organizing protein that folds chromatin by tethering specific genomic elements, known as base-unpairing regions (BURs), which have a high propensity for unwinding^10^, to the SATB1 nuclear architecture^11^. Through this activity, SATB1 establishes cell type-specific three-dimensional chromatin architecture and globally reprograms gene expression, thereby driving phenotypic plasticity in host cells^12^. These functions contribute to postnatal T-cell development and function^13–18^, differentiation of multiple cell lineages^19–25^, and acquisition of metastatic phenotypes in cancer^1,2^.

Using human breast cancer cell lines in mouse models we previously demonstrated that SATB1 is a key determinant of highly invasive tumor cell phenotypes^1,2^. SATB1 protein expression in the nuclei of breast tumor cells is associated with poor prognosis of patients with breast cancer, independent of lymph node status (P<0.001). Importantly, SATB1 protein is absent in normal breast tissue but is expressed in metastatic breast cancers, including many cases of triple-negative breast cancer (TNBC). It is of interest to note that although normal breast tissue does not express SATB1 protein^1^, it does uniformly transcribe medium to high levels of mRNA for the *SATB1* gene based on The Genotype-Tissue Expression (GTEx). While nearly 1,000 genes show SATB1-dependent changes in expression as breast cancer cells acquire metastatic properties, no studies have focused on identifying actionable downstream target genes of SATB1. Because SATB1 itself is also transcribed in normal breast tissue, SATB1 target genes may instead serve as more specific RNA-based markers for assessing the risk of breast cancer metastasis. The information obtained with breast cancer may be applicable to other cancer types because SATB1’s high prognostic significance have been confirmed in approximately 20 different types of epithelial cancers, including pancreatic cancer, prostate cancer, and head and neck squamous carcinoma (HNSCC)^1–9^.

Given that SATB1 serves as a critical driver of metastasis when expressed in human epithelial cancers, we focused on genes involved in epithelial-to-mesenchymal transition (EMT) as potential target genes of SATB1. EMT is a crucial process in embryonic development, during which non-motile epithelial cells acquire the ability to migrate, becoming motile mesenchymal stem cells. When aberrantly activated in cancer, EMT-promoting genes enhance the invasive behavior of carcinoma cells and plays a fundamental role in the metastatic progression of cancer, facilitating the migration of tumor cells from the primary site to distant locations where they can grow^26,27^. Additionally, in cancer EMT enhances cancer stem cell activity, increases resistance to chemotherapy, and promotes immune evasion^28–30^. EMT is a complex and dynamic process involving many genes with diverse biological functions that both respond to and remodel the tumor extracellular environment and tumor-cell properties^31^. These EMT genes may collectively contribute to the key mechanisms of EMT underlying cancer progression, metastasis, and resistance to treatment. There are many EMT-promoting genes with distinct cellular roles, but it remains unknown whether any specific subset of EMT genes is crucial for cancer metastasis. We hypothesized that SATB1 activates a critical subset of EMT genes during breast cancer progression toward metastasis. To test this hypothesis, we generated a new gene cohort consisting of 300 EMT-promoting genes, incorporating newly identified EMT-promoting genes from recent literature, together with 39 additional genes, including 27 EMT-inhibitory genes and 12 metastasis-suppressor genes. We then examined a total of 339 genes involved in EMT-related processes in cancer to determine whether their expression is dependent on SATB1.

To identify SATB1-regulated EMT genes, we used a modified version of the non-malignant immortalized breast epithelial cell line MCF10A, designated MCF10A-1. This variant cell line has been well characterized and develops highly invasive and metastatic phenotypes following SATB1 transduction^1^. In contrast to normal breast tissue, *SATB1* mRNA is essentially undetectable in parental MCF10A cells, which also do not express SATB1 protein^1^. Moreover, parental MCF10A cells do not respond to ectopic SATB1 expression, whereas MCF10A-1 cells are permissive to SATB1-induced phenotypic changes. These changes include altered cell morphology, increased invasive activity in vitro, and enhanced tumor growth at xenografted sites and in the lungs (experimental metastasis) of immunocompromised mice^2^.

MCF10A-1 cells were generated after repeated passages in culture, which exhibited significant alterations in cell cycle-related and checkpoint genes, in particular major reduction of the early DNA damage-response kinases Ataxia telangiectasia mutated (ATM) and Rad3 related (ATR)^2^. There is no difference in genomic integrity between MCF10A and MCF10A-1 cells. ATM is crucial for the mitotic checkpoint in the cell cycle^32^, and MCF10-1A cells with low ATM levels show an impaired G2/M checkpoint^2^. Stable ATM knockdown in parental MCF10A cells, as well as in the non-malignant breast epithelial cell lines 184A1 and 184B5, creates a SATB1-permissive state that enables SATB1-mediated invasive activity *in vitro*. To date, among non-tumorigenic breast epithelial cell lines, only MCF10A-1 cells have been confirmed *in vivo* to exhibit SATB1-induced tumorigenic and metastatic properties. Therefore, MCF10A-1 cells provide a valuable model for studying the transitional state of premalignant cells poised for malignant transformation and metastatic progression in a SATB1-dependent manner. Accordingly, the MCF10A-1 cell model is uniquely suited for examining SATB1-mediated activation of EMT genes during malignant transformation. Notably, low ATM expression is associated with poor prognosis in multiple cancers, including breast cancer^33,34^, further supporting the notion that MCF10A-1 cells have undergone a transition resembling the natural progression toward malignancy. Using two genome-wide transcription methodologies, RNA-seq and GRO-seq (Global Run-On sequencing), we explored SATB1-dependent gene expression among the discrete 339 gene set involved in EMT. This led to the identification of 98 EMT-promoting genes that SATB1 upregulates in MCF10A-1 cells.

In this study, we also tested the hypothesis that a distinct etiology, such as the environmental chemical carcinogen benzo[a]pyrene (B[a]P), activates a similar subset of EMT genes targeted by SATB1. If SATB1-upregulated EMT-promoting genes play particularly important roles in breast cancer, then exposure to B[a]P would also activate a significantly overlapping set of EMT genes. B[a]P is one of the best characterized components of polycyclic aromatic hydrocarbon (PAH), is produced during incomplete combustion of organic material such as fossil fuels and wood, and it is found in automobile exhaust fumes and tobacco smoke^35,36^. Carcinogenicity of B[a]P has been well demonstrated in many experimental animal models^37,38^. Direct DNA damage by forming DNA adduct formations through a metabolite, B[a]P diol epoxide (BPDE), is considered as a crucial step in the initiation of carcinogenesis^39,40^. B[a]P has been shown to increase the risk of breast cancer, sarcomas, liver and lung tumours^41,42^. The risk of breast cancer is associated with long-term cumulative exposure of B[a]P, shown by the result of a large cohort of patients and controls (5222 cases each) with 21 years of follow-up period^42^. Using human breast epithelial and breast cancer cell lines, both *in vitro* and in mouse models *in vivo*, B[a]P has been shown to induce tumor growth, invasive activity, and experimental metastasis^43–46^. Short-term exposure of MCF10A-1 cells to B[a]P upregulated 72 EMT genes, showing a statistically significant 50% overlap with EMT genes upregulated by SATB1. These findings support the notion that a specific subset of EMT genes plays a critical role in malignancy despite distinct oncogenic etiologies. Unlike all other non-SATB1-regulated EMT genes, SATB1-activated EMT genes were significantly enriched in the BL2 subset of TNBC, which is associated with the poorest prognosis among TNBC subgroups^47^, highlighting their translational significance. This specific set of identified SATB1-dependent EMT genes is not merely a random collection; it likely represents the key molecular events involved in the malignant transformation of breast epithelial cells as they acquire metastatic properties.

## Results

Using the SATB1-transduced MCF10A-1 cell line, which acquires malignant phenotypes, we analyzed SATB1-dependent transcriptional changes associated with malignant transformation. We compared transcripts from MCF10A-1 cells transfected with either a vector control or a SATB1 expression construct^2^ using parallel RNA-seq and GRO-seq analyses. SATB1-dependent differential gene expression results are presented in Supplementary Table 1. Whereas RNA-seq measures steady-state RNA levels, GRO-seq captures nascent transcripts synthesized by RNA polymerase II (Pol II) in nuclei^48^, enabling identification of genes more likely to be directly regulated by SATB1. Because GRO-seq is also well suited for detecting rapid transcriptional responses to external stimuli, we primarily used GRO-seq to compare nascent transcripts induced by the chemical carcinogen benzo[a]pyrene (B[a]P) with those activated by SATB1, as described below.

### Functional Annotation of SATB1-Dependent Genes

By the DAVID functional annotation tool^49^, we first examined data from RNA-seq to identify functional gene clusters that are highly enriched in SATB1-dependent genes. A total of 585 genes (identified in DAVID) that were upregulated [≥ 2-fold, adjusted p value (p_adj_) ≤ 0.05] in SATB1-expressing MCF10A-1 cells, compared to control MCF10A-1 cells, were found to be significantly enriched in functionally related gene groups, such as animal organ development, extracellular matrix (ECM), regulation of cell migration, cell communication, DNA-binding transcription factor activity (Figure 1A). Because SATB1 has multiple roles in T cells, we also reviewed genes under the category “regulation of T cell activation”. Despite a moderate enrichment, the transcript levels of the programmed death-ligand 1 (*PD-L1/CD 274*) and PDCD1LG2 (*PD-L2/CD272*) were upregulated by 2-fold (p_adj_ =2.23x10^-9^) or 3-fold (p_adj_ =1.04x10^-7^), respectively, depending on SATB1. PD-L1 and PD-L2 bind to the PD-1 receptor on activated T cells, effectively hindering the immune system from attacking cancer cells and allowing them to evade destruction^50–52^.

**Figure 1.**
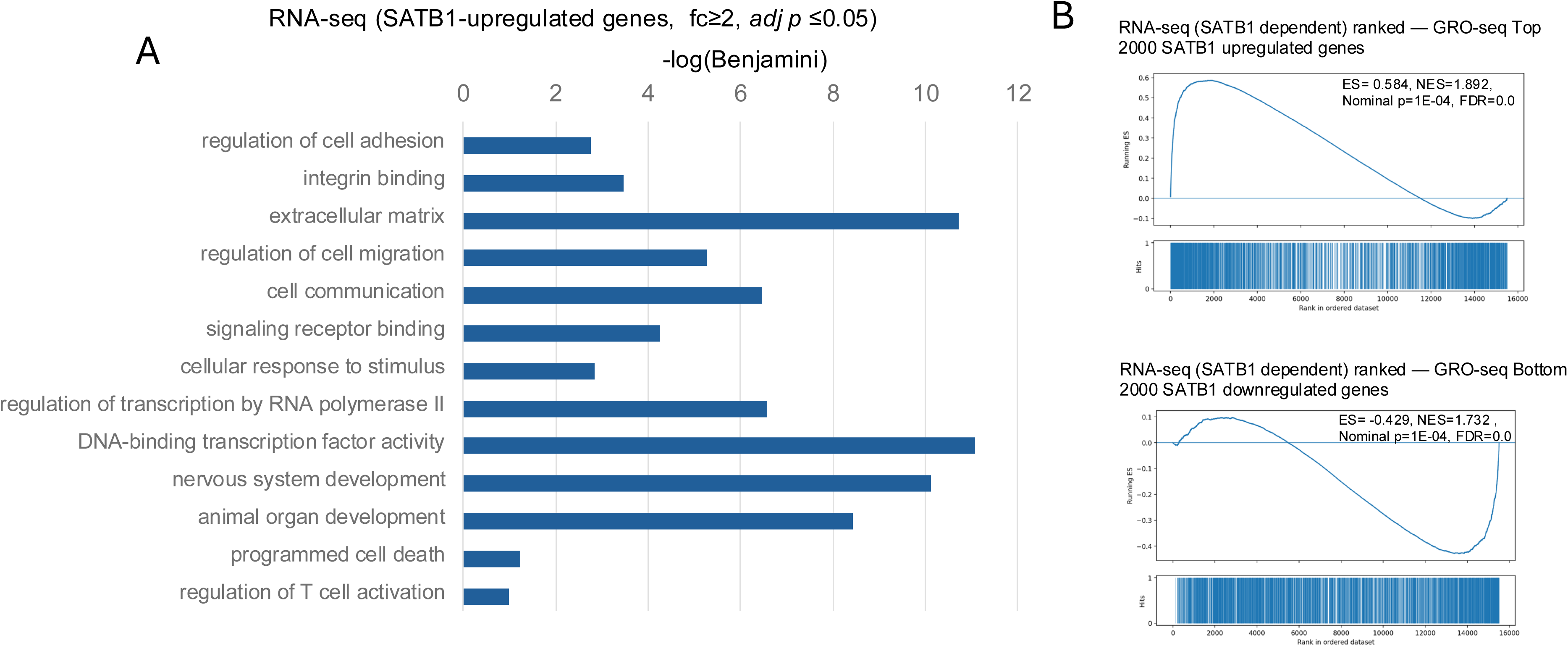
Analysis of SATB1-regulated genes. **A)** DAVID functional annotation analysis of 585 SATB1-upregulated genes (fc ≥2, p_adj_ ≤0.05) in MCF10A-1 compared with control non SATB1-expressing cells determined by RNA-seq. X-axis [-log (Benjamini)] indicates the statistical significance of each gene group. **B)** Gene Set Enrichment Analysis (GSEA) using RNA-seq genes ranked by log2 fold change and tested gene sets defined by the top and bottom 2,000 GRO-seq–SATB1-regulated genes. GRO-seq upregulated genes are enriched among RNA-seq upregulated genes (top panel), while GRO-seq downregulated genes are enriched among RNA-seq downregulated genes (bottom panel), demonstrating strong directional concordance between nascent transcriptional regulation and steady-state RNA expression. Enrichment scores (ES) were calculated using the Broad Institute preranked GSEA algorithm (weight p = 1) with 10,000 permutations, and normalized enrichment scores (NES) were derived using same-sign null distributions. Nominal p-values and false discovery rates (FDR) were computed according to Broad Institute methodology. Positive NES indicates enrichment toward the top of the RNA-seq ranked list, whereas negative NES indicates enrichment toward the bottom. Gene-set hit positions are indicated by thin vertical tick marks in the separate “Hits” panel below.

Many functional gene groups analyzed using RNA-seq data (Figure 1A) were commonly enriched among SATB1-dependent genes identified by GRO-seq (Supplementary Table 2). To compare the two datasets at the individual gene level, we analyzed a comparable number of GRO-seq–derived genes (654 genes, identified by DAVID) using a threshold of fold change (fc) ≥ 1.7 and nominal p ≤ 0.05. We intentionally used nominal p values for GRO-seq to test whether a similar set of SATB1-dependent genes could be identified by GRO-seq as by RNA-seq (filtered with p_adj_ ≤ 0.05). In SATB1-expressing cells, GRO-seq measurements showed greater variability across replicates than RNA-seq. This variability may reflect enhanced dynamics in RNA polymerase activity uniquely associated with SATB1 expressing cells, leading to increased cell-to-cell variation in nascent transcript levels (see Discussion). Nevertheless, under the filtration thresholds employed, GRO-seq captured many key SATB1-dependent genes consistent with RNA-seq results (Supplementary Table 2).

To directly assess concordance between nascent transcripts detected by GRO-seq and steady-state expression measured by RNA-seq, we performed a Gene Set Enrichment Analysis (GSEA) test that does not require prior filtration by p values. Using RNA-seq genes ranked by log2 fold change depending on SATB1, gene sets defined by the top 2,000 GRO-seq upregulated genes and the bottom 2,000 GRO-seq downregulated genes were significantly enriched at the top end of the RNA-seq ranked list (SATB1-upregulated genes) or at the bottom end of the RNA-seq ranked list (downregulated genes) (p=0.0001, FDR=0.000 in both cases). This demonstrates strong concordance and directional agreement between GRO-seq–defined transcriptional regulation and RNA-seq–defined expression changes (Figure 1B). Furthermore, the top 100 SATB1-dependent genes (based on fold change) in RNA-seq and GRO-seq, analyzed by Reactome, revealed that 26 of the 35 highest enrichment pathways were common to both RNA-seq and GRO-seq, highlighting their SATB1 dependence (Supplementary Table 1). These common functional categories and pathways dependent on SATB1 are highly relevant to tumor growth and metastasis.

### Identification of 98 EMT genes regulated by SATB1

Next, we investigated the impact of SATB1 on genes known to contribute to the invasive activity of cancer cells. We specifically focused on genes involved in EMT, a process that enables tumor cells to acquire the invasive capabilities required for metastasis^27,28,31^. Because SATB1 transduction in MCF10A-1 cells induces invasive activity both *in vitro* and *in vivo*, including lung colonization following tail-vein injection into mice^2^, EMT genes are likely to represent key targets of SATB1. We further hypothesized that SATB1 preferentially upregulates a specific subset of EMT-promoting genes. To comprehensively analyze the impact of SATB1 on EMT-related genes beyond a limited number of canonical EMT genes, we compiled an EMT gene set consisting of 339 genes. This gene set was generated by reviewing genes associated with two Gene Ontology (GO) terms related to EMT regulation (GO:0010717 and GO:0001837), incorporating genes from the Qiagen GeneGlobe database, and adding numerous EMT-promoting genes identified in recent literature. The EMT gene set comprised 300 EMT-promoting genes, 27 EMT-inhibitory genes, and 12 metastasis suppressor genes^53,54^ (Supplementary Table 3). Most EMT-promoting genes included in this set have been confirmed to facilitate EMT in breast cancer.

Preranked GSEA revealed strong concordant enrichment of the 300 EMT-promoting gene set in both RNA-seq- and GRO-seq-ranked gene lists (p=0.0001, FDR=0.000 for both) (Figure 2A). This indicates that EMT-associated genes are preferentially upregulated at both the nascent transcript and steady-state expression levels. Among the top 50 genes from RNA-seq that confer strong SATB1 dependency, fold change ranged from fc=2178 (p=8.2x10^-19^) for *CXCL5* to fc=37 (p=1.2x10^-20^) for *SPARC*. Of these, 20 genes were involved in EMT processes based on our EMT gene list. This finding was consistent for GRO-seq as well, with 21 of the 50 top SATB1-dependent genes also classified as EMT-promoting genes (Supplementary Table 1).

**Figure 2.**
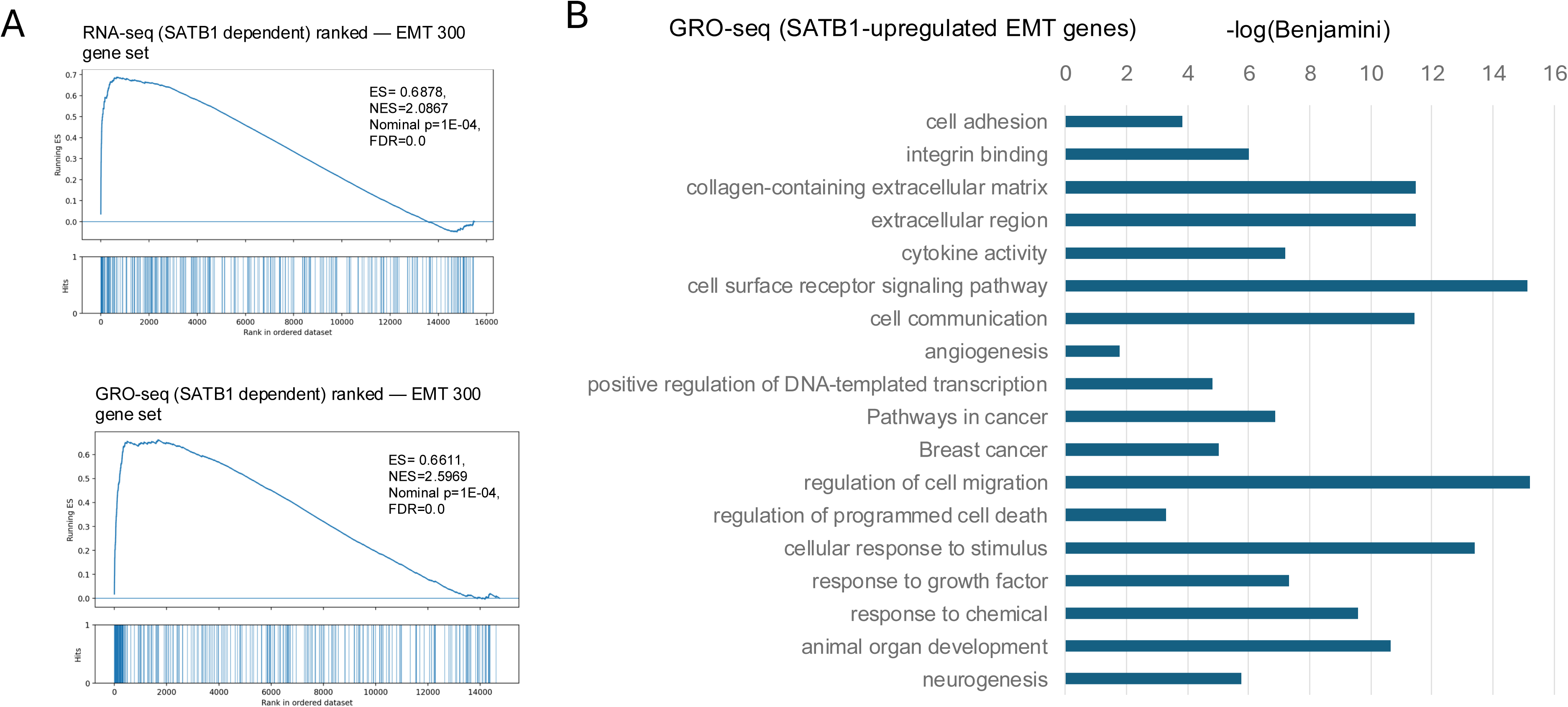
Analysis of SATB1-regulated EMT-promoting genes**. A)** GSEA was performed independently using genes ranked by log2 fold change (log2FC) from RNA-seq and GRO-seq datasets. A curated gene set comprising 300 EMT-promoting genes was tested for enrichment. Enrichment scores (ES), normalized enrichment scores (NES), nominal p-values and false discovery rates (FDR) are indicated. Gene-set hit positions are indicated by thin vertical tick marks in a separate “Hits” panel below. **B)** DAVID gene enrichment analysis of 98 SATB1-upregulated EMT-promoting genes in SATB1- transduced MCF10A-1 compared with control cells (fc > 1.2, p < 0.07) determined by GRO-seq. X-axis [-log (Benjamini)] indicates the statistical significance of each gene group.

From the curated list of 300 EMT-promoting genes, we identified 98 genes that are upregulated by SATB1 based on GRO-seq data (Table 1). This group of 98 genes includes 86 genes (fc ≥ 1.2, p ≤ 0.05) and an additional 12 genes (fc ≥ 1.2, 0.05 p ≤ 0.07). EMT-promoting genes are found to be significantly enriched among SATB1-upregulated genes relative to all other genes (including SATB1-downregulated genes and all non-SATB1-dependent genes) with Odds Ratio (OR)= 5.53 and two-sided Fisher’s exact test: p = 1 × 10⁻³⁰. The functions of these SATB1-upregulated EMT-promoting genes and their relevance to cancer are summarized in Supplementary Table 4.

**Table 1.** 98 SATB1-upregulated EMT-promoting genes identified from GRO-seq.

| Gene ID | fold change | Gene ID | fold change | Gene ID | fold change | Gene ID | fold change |
| --- | --- | --- | --- | --- | --- | --- | --- |
| CXCL5 | 25.69 | FBXO32 | 5.33 | CREB3L1 | 3.46 | TIMP1 | 1.94 |
| COL1A2 | 24.30 | CEMIP | 5.25 | SULF1 | 3.42 | VEGFA | 1.88 |
| IGFBP3 | 23.78 | ASIC3 | 5.16 | BNC2 | 3.38 | WNT2B | 1.87 |
| SPARC | 18.18 | FOXG1 | 4.94 | WNT5B | 3.37 | ID2 | 1.87 |
| CDKN2A | 12.34 | CCDC102B | 4.83 | HEY1 | 3.31 | SMAD7 | 1.80 |
| STC1 | 11.79 | PLIN2 | 4.73 | EGFL7 | 3.23 | FN1 | 1.80 |
| FABP4 | 10.71 | CD68 | 4.65 | MMP3 | 3.21 | ITGA2 | 1.79 |
| ADAMTS5 | 10.70 | NRSN2 | 4.60 | TCF7 | 3.14 | JAG1 | 1.77 |
| SHISA2 | 10.53 | HBEGF | 4.47 | GFPT2 | 3.09 | IL1B | 1.74 |
| THY1 | 10.24 | AKR1C4 | 4.45 | CCN4 | 3.07 | HIF1A | 1.66 |
| IL13RA2 | 10.11 | ABCA1 | 4.42 | FOSB | 3.06 | RELB | 1.66 |
| COL3A1 | 9.86 | TMEM158 | 4.37 | LIN7A | 3.04 | FOXC1 | 1.65 |
| THBS2 | 8.74 | SPP1 | 4.28 | PSG9 | 3.02 | FERMT2 | 1.57 |
| ADAM28 | 8.44 | NGF | 4.25 | BMP7 | 3.01 | JAK1 | 1.56 |
| ZFP3 | 8.42 | SMAD6 | 4.16 | LIF | 2.93 | APLF | 1.56 |
| NFE2 | 7.16 | MMP9 | 4.12 | FGF23 | 2.90 | FZD6 | 1.55 |
| CHAC1 | 7.05 | COL5A2 | 4.09 | FZD2 | 2.89 | APH1B | 1.54 |
| SOX5 | 6.49 | LAPTM5 | 4.06 | PSENEN | 2.87 | PDPN | 1.50 |
| PTX3 | 6.10 | NUPR1 | 3.90 | ADAM19 | 2.72 | SOS2 | 1.33 |
| TNFSF10 | 5.99 | USP27X | 3.82 | EGR1 | 2.68 | NCSTN | 1.31 |
| UCHL1 | 5.87 | GDF15 | 3.77 | GCNT2 | 2.54 | PARD6B | 1.30 |
| ARHGEF19 | 5.69 | IL13 | 3.76 | COL1A1 | 2.52 | MET | 1.28 |
| POSTN | 5.64 | SMO | 3.61 | MAPK3 | 2.14 | ADAM17 | 1.27 |
| COL5A1 | 5.63 | NDUFA4L2 | 3.53 | NFKB2 | 2.01 |  |  |
| IL33 | 5.57 | GSDMB | 3.47 | PHLDB2 | 1.98 |  |  |

Genes were selected by fold change $\geq 1.2$ and $p \leq 0.05$ , except for genes (blue) by $p \leq 0.07$ .

Among 27 EMT-inhibiting genes, *OVOL2* (fc=0.60, p=0.04), *CDH1* (fc-0,49, p=0.01, *EMP2* (0.48, p=0.005), and *DKK1* (fc=0.25, p=0.0006) to be downregulated by SATB1 by GRO-seq and verified by RNA-seq data. In addition, among the 12 metastasis suppressor genes, GRO-seq detected *ARHGDIB* (fc=0.50, p=0.0059) and *CLDN4(Claudin-4*) (fc=0.56, p=0.06) to be significantly downregulated in SATB1-expressing MCF10A-1 cells. From RNA-seq data (not from GRO-seq), *TXNIP* (fc=0.57, p=2.65x10^-17^), *GSN* (*Gelsolin*) (fc=0.70, p=1.35x10^-10^) and *MAP2K6* (fc=0.68, p= 0.003) werec significantly downregulated by SATB1. Other well-known metastasis suppressor genes, i.e. *BRMS1* and *KiSS1* remained essentially unchanged by SATB1 in MCF10A-1.

### SATB1 upregulates EMT-promoting genes with ECM functions and Transcription

EMT involves a heterogeneous set of genes with mechanistically distinct functions, including ECM remodeling, cytoskeletal reorganization, transcriptional reprogramming, survival signaling, and metabolic adaptation. Despite their diverse functions, these genes collectively drive invasive and metastatic phenotypes^26–31^. We analyzed functional categories of 98 SATB1-upregulated EMT-promoting genes using DAVID (Figure 2B; Table 2; Supplementary Table 5). Twenty-one genes were associated with cancer pathways, including 11 specifically linked to breast cancer. Representative genes from selected functional categories are described below.

**Table 2.**
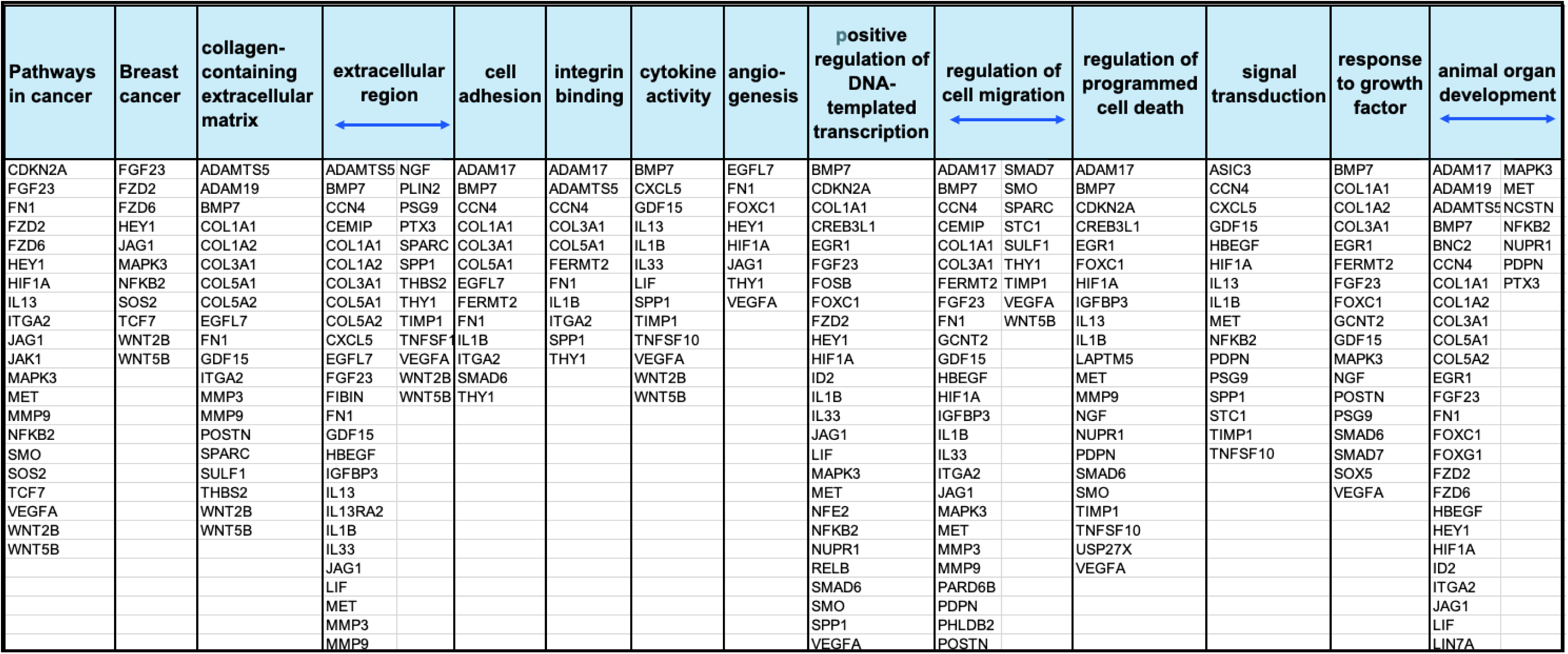
SATB1-upregulated EMT-promoting genes (GRO-seq.

### ECM components

ECM is a three-dimensional macromolecular network that surrounds cells and is essential for maintaining the structure and function of all tissues. It consists of structural proteins (such as collagen, elastin, and fibronectin), proteoglycans, and glycoproteins, which together form a large fibrillar structure. The most significantly enriched category among SATB1-dependent EMT genes comprised ECM-associated proteins, particularly collagen-containing ECM. SATB1 strongly induced five collagen genes (*COL1A1, COL1A2, COL3A1, COL5A1*, and *COL5A2*), with *COL1A2* upregulated 24.3-fold (p=0.031), second only to *CXCL5.* Additional collagen genes were also robustly induced (*COL3A1, COL5A1, COL1A1*). Collagen-rich matrices are key determinants of the tumor microenvironment and are associated with hypoxia, chemoresistance, invasion, and metastasis. SATB1-dependent collagen genes (*COL1A1, COL1A2, COL5A1*) are frequently overexpressed in gastric, breast, and colorectal cancers and correlate with tumor proliferation, metastasis, and drug resistance^55,56^. In breast cancer, elevated *COL1A2* and *COL3A1* expression, particularly in TNBC, is associated with reduced overall and recurrence-free survival ^57–59^.

### Chemokines, cytokines, and growth factors

These factors are considered functional components of ECM because ECM acts as a bioactive scaffold that binds and controls the availability of these signaling proteins. SATB1 induced multiple secreted EMT-promoting factors, including a chemokine, *CXCL5*, cytokines *(IL13, IL1B, IL33, GDF15, LIF, SPP1, TNFSF10*), and growth factors (BMP7, NGF, VEGFA). *CXCL5* was the most strongly upregulated SATB1 target detected in both GRO-seq and RNA-seq data. *CXCL5* has been implicated in tumor cell proliferation, angiogenesis, metastatic migration, and reactivation of dormant breast cancer cells that disseminate to the bone^60–63^. SATB1 also increased expression of CXCR2 (detected by RNA-seq), the CXCL5 receptor, reinforcing a transcriptional program that enhances CXCL5-driven metastasis.

### Cell–ECM communication and ECM remodeling

SATB1 upregulates numerous genes involved in cell–ECM communication that are critical for ECM remodeling, through which tumors create a microenvironment that promotes tumorigenesis and metastasis^64^. These include *THBS2* ^65,66^, *POSTN (Periostin)*^67^, *PDPN (Podoplanin)*^68^, *FN1*^69,70^, and *VEGFA*^71,72^. All these proteins when dysregulated play roles in remodeling the tumor microenvironment (TME), promoting angiogenesis (VEGFA being a key driver), facilitating cell migration and invasion (metastasis), and increasing matrix stiffness, a hallmark of aggressive tumors associated with enhanced mechanotransduction signaling and poor patient prognosis^73^. SATB1 also upregulates SPARC, which exerts significant effects on tissue architecture^74^and has similar activities (angiogenesis, migration, proliferation and survival). However, *SPARC* has been reported to either promote or suppress cancer progression depending on the cancer type and TME^75^. In TNBC, *SPARC* overexpression correlates with disease recurrence and poor overall survival in patients^76^.

SATB1 also induced genes encoding ECM-remodeling enzymes, including metalloproteinases (*MMP3, MMP9*)^77^, *ADAM* family members (*ADAM17, ADAM19, ADAM28*), and *ADAMTS5*^78,79^, and the hyaluronidase CEMIP^80^. These enzymes remodel ECM components, regulate shedding of cell-surface proteins and matrix-bound growth factors, and remove physical barriers to migration, thereby creating a pro-invasive tumor microenvironment^81^. In addition, SATB1 upregulated genes encoding enzymes involved in post-translational modification, including the E3 ubiquitin ligase *FBXO32* and the deubiquitinase *UCHL1*. *FBXO32* is elevated in metastatic cancers, and its tumor growth and metastasis-promoting functions have been demonstrated *in vivo*^82^. UCHL1, a novel target in breast cancer^83–85^, enhances TGF-β signaling-induced metastasis by inhibiting the degradation of the TGF-β type I receptor and SMAD2.

### Cancer-relevant signaling pathways

By KEGG and Wiki pathway analyses, 59 of the 98 SATB1-upregulated EMT genes were found to participate in one or more cancer-relevant signaling pathways, including pathways in cancer and breast cancer, with Benjamini-Hochberg=2.00x10^-9^ or 1.90x10^-7^, respectively, as well as PI3K–Akt, Wnt, Notch, Hippo, TNF, MAPK, TGF-β, mTOR, and FoxO signaling (Supplementary Table 5). Importantly, multiple SATB1-dependent EMT genes converged on Wnt signaling, including *WNT2B, WNT5B, FZD2, FZD6, and TCF7*. WNT2B and WNT5B are secreted glycoproteins that bind to cell surface receptors, initiating intracellular signaling pathways that control gene transcription. Frizzled receptors (FZDs; FZD2 and FZD6) are the primary receptors for Wnt ligands and are essential for Wnt signaling. TCF7 proteins are key downstream effectors of the Wnt signaling pathway, regulating gene expression. In addition, *WISP1/CCN4*, a Wnt target gene upregulated by SATB1, promotes EMT, invasion, and metastasis in breast cancer^86–89^.

SATB1 also induced genes encoding several components of the Notch signaling pathway (Supplementary Table 5), including *JAG1* for the Jagged 1 protein, a Notch ligand strongly associated with breast cancer metastasis^90^. Additional SATB1-regulated genes encode proteolytic components required for NOTCH receptor activation, including *ADAM17* and multiple γ-secretase complex components (*APH1B, PSENEN, NCSTN*). SATB1 further upregulated Smoothened (SMO), a core Hedgehog pathway effector implicated in breast cancer progression, metastasis, and cancer stem cell maintenance^91^. These SATB1-regulated pathways—Wnt, Notch, Hedgehog, and TGF-β—are highly interconnected, and their coordinated dysregulation promotes tumor growth, invasion, metastasis, and treatment resistance^92,93^.

Beyond canonical oncogenic pathways, SATB1 upregulated genes involved in cell–cell and cell–ECM signaling, including *THY1* and *MET*. THY1 contributes to cancer stemness and invasion through pathways such as Wnt/β-catenin and AMPK/mTOR, whereas MET encodes a proto-oncogenic receptor tyrosine kinase that promotes tumor growth, invasion, metastasis, and chemoresistance^94–96^.

### Transcriptional factors and canonical EMT markers

Among SATB1-upregulated EMT genes (Table 2; Supplementary Table 4), several higher-order regulators of EMT-associated transcriptional programs were prominent. These included *HIF1A*, a central mediator of hypoxia-driven transcriptional reprogramming^97,98^; *HEY1*, a Notch pathway effector^99^; *TCF7,* a Wnt/β-catenin transcriptional regulator^100^; *RELB,* a component of non-canonical NF-κB signaling^101^; and *FOXC1*, an EMT-associated transcription factor linked to aggressive tumor phenotypes^102,103^. The coordinated induction of these regulators indicates activation of multiple convergent transcriptional circuits governing EMT, stemness, inflammatory signaling, and tumor progression. Consistent with EMT induction, SATB1 repressed *CDH1 (E-cadherin*) detected by both GRO-seq and RNA-seq. However, canonical EMT transcription factor genes such as *SNAI1/SNAI2, ZEB1/ZEB2*, and *TWIST1/TWIST2* were not activated by SATB1 (by GRO-seq). Only in RNA-seq, *SNAI1/SNAI2* and *ZEB1* were marginally upregulated (fc=1.2 to 1.6).

These findings collectively show that SATB1 activates key genes involved in ECM remodeling, structural and functional components of the ECM, oncogenic signaling, and master transcriptional regulators. The coordinated activation of these genes with these diverse but complementary functions is expected to effectively promote tumor progression and metastasis.

### SATB1-regulated EMT genes coordinate immune evasion and angiogenesis

The coordinated activation of angiogenesis and immune evasion is increasingly recognized as a central mechanism underlying tumor progression, metastatic dissemination, and therapeutic resistance. Because EMT also involves immune modulation and vascular remodeling in addition to invasion and motility^29^, we asked whether SATB1-regulated EMT genes contribute to these tumor-promoting processes. Angiogenesis and immune evasion frequently co-occur in tumors because they are driven by shared upstream pressures, particularly hypoxia and chronic inflammation, and coordinated through regulatory hubs such as HIF1A, STAT3, NF-κB, and TGF-β. EMT programs integrate these signals to remodel both the tumor vasculature and the immune microenvironment, linking metastatic competence with immune escape and vascular dysregulation^29,104^.

The ECM remodeling factors described above (i.e. POSTN, THBS2, FN1, PDPN, VEGFA, MMP9, and ADAM17) promote angiogenesis and immune evasion through coordinated remodeling of the extracellular, vascular, and inflammatory architecture of the tumor microenvironment (TME). VEGFA, a canonical angiogenic factor, drives endothelial proliferation and vascular permeability. Beyond its role in angiogenesis, VEGF functionally links angiogenesis and immune evasion by generating abnormal vasculature that restricts immune-cell infiltration while promoting immunosuppressive signaling, dendritic-cell dysfunction, recruitment of suppressive myeloid populations, and expansion of regulatory T cells^105^.

Several of these factors, including FN1, POSTN, and THBS2, are frequently enriched in cancer-associated fibroblasts (CAFs) and contribute to fibrotic stromal remodeling. However, they can also be produced by EMT-like tumor cells generated through SATB1 activation. FN1 is a canonical mesenchymal marker and a hallmark of EMT when expressed by tumor cells. These tumor-derived stromal proteins may similarly promote immune exclusion and angiogenesis. Indeed, tumor cells undergoing EMT actively remodel the TME and, in some contexts, acquire CAF-like functions^104^.

Among the ECM-remodeling proteases, MMP9 and ADAM17 further integrate angiogenic and immunosuppressive programs. MMP9 mobilizes VEGF sequestered within the ECM, thereby coupling matrix remodeling to angiogenesis and immune suppression^106^. ADAM17 has been implicated in tumor immune evasion through shedding of PD-L1 and ligands involved in NK-cell-mediated immune surveillance, while also promoting a pro-tumor inflammatory TME^107,108^. In parallel, SATB1-induced HIF1A transcriptionally activates angiogenic programs, including VEGF signaling, thereby enhancing tumor angiogenesis and reinforcing an immunosuppressive TME^97^. MET similarly promotes neovascularization, invasive growth, and immune evasion^95,96^.

The SATB1 program also incorporates genes associated with developmental immune tolerance pathways. Notably, PSG9 (pregnancy-specific glycoprotein 9) promotes tumor growth, angiogenesis, and metastasis while enhancing FoxP3⁺ regulatory T-cell differentiation and function, exemplifying tumor co-option of developmental immune tolerance (“placental mimicry”) programs^109,110^.

Within cytokine and chemokine networks, SATB1-regulated EMT further amplifies vascular–immune coupling. CXCL5, a CXCR2-binding chemokine, promotes endothelial activation and angiogenesis while recruiting immunosuppressive myeloid-derived suppressor cells (MDSCs) to the TME^60^. IL33, a context-dependent alarmin, enhances vascular activation and expansion while driving immunosuppressive inflammatory programs through myeloid and stromal signaling networks^60^. SPP1 (osteopontin) functions as a secreted immunomodulatory factor that supports endothelial migration and vessel formation while promoting recruitment and polarization of tumor-associated macrophages toward immunosuppressive phenotypes^111^.

SATB1-regulated EMT genes collectively orchestrate microenvironmental remodeling through hypoxia-responsive pathways, inflammatory mediators, extracellular matrix remodeling, and myeloid cell recruitment. This SATB1-regulated EMT program functionally links two interdependent processes—angiogenesis and immune evasion—that work together to enhance metastatic competence.

### Enrichment of SATB1-Activated EMT Genes Primarily in the BL2 subtype of TNBC

TNBC lacks expression of estrogen receptor (ER), progesterone receptor (PR), and human epidermal growth factor receptor 2 (HER2) and represents approximately 10–20% of all breast cancers^112,113^. TNBC remains one of the most aggressive type of breast cancer and is marked by high recurrence rates, early metastasis, limited treatment options, and poor overall survival. Importantly, TNBC is a highly heterogeneous disease: while many patients experience rapid disease progression and metastasis, others have more indolent disease courses. Among patients with TNBC who received no treatment (no surgery, chemotherapy, or radiation), approximately 19% survived longer than 10 years^114^. Based on analysis of transcriptional profiling and gene pathways, TNBCs are classified into four subtypes**: (**Basal-Like 1 (BL1), Basal-Like 2 (BL2), Mesenchymal (M), and Luminal Androgen Receptor (LAR) by Lehmann et al^115^. These subtypes differ markedly in biology, therapeutic vulnerabilities, and outcomes. The BL2 subtype is characterized by high chemoresistance and clinical aggression, leading to early recurrence and exhibiting the lowest pathological complete response (pCR) rates, often below 10%^116,117^. This subtype has a poorer prognosis compared to other TNBC subtypes.

To evaluate the clinical relevance of the 98 SATB1-activated EMT genes identified in MCF10A-1 cells, we examined their distribution across TNBC subtypes. SATB1-dependent EMT genes showed a striking and highly significant enrichment in the BL2 subtype (49 of 98 genes; Fisher’s exact test, p = 4.65 × 10⁻⁹) (Figure 3A and B). A significant association was also observed with the mesenchymal (M) subtype (28 of 98 genes; Fisher’s exact test, p = 0.0035), which is characterized by invasive, migratory features and EMT gene expression. In contrast, no significant enrichment was detected in BL1 or LAR tumors (Fisher’s exact test, p = 0.34 and 0.40, respectively). Notably, BL1, the subtype with the most favorable prognosis and highest chemotherapy response rates, exhibits the lowest representation of SATB1-activated EMT genes, with only 18 of 98 genes present (Figure 3A and 3B).

**Figure 3.**
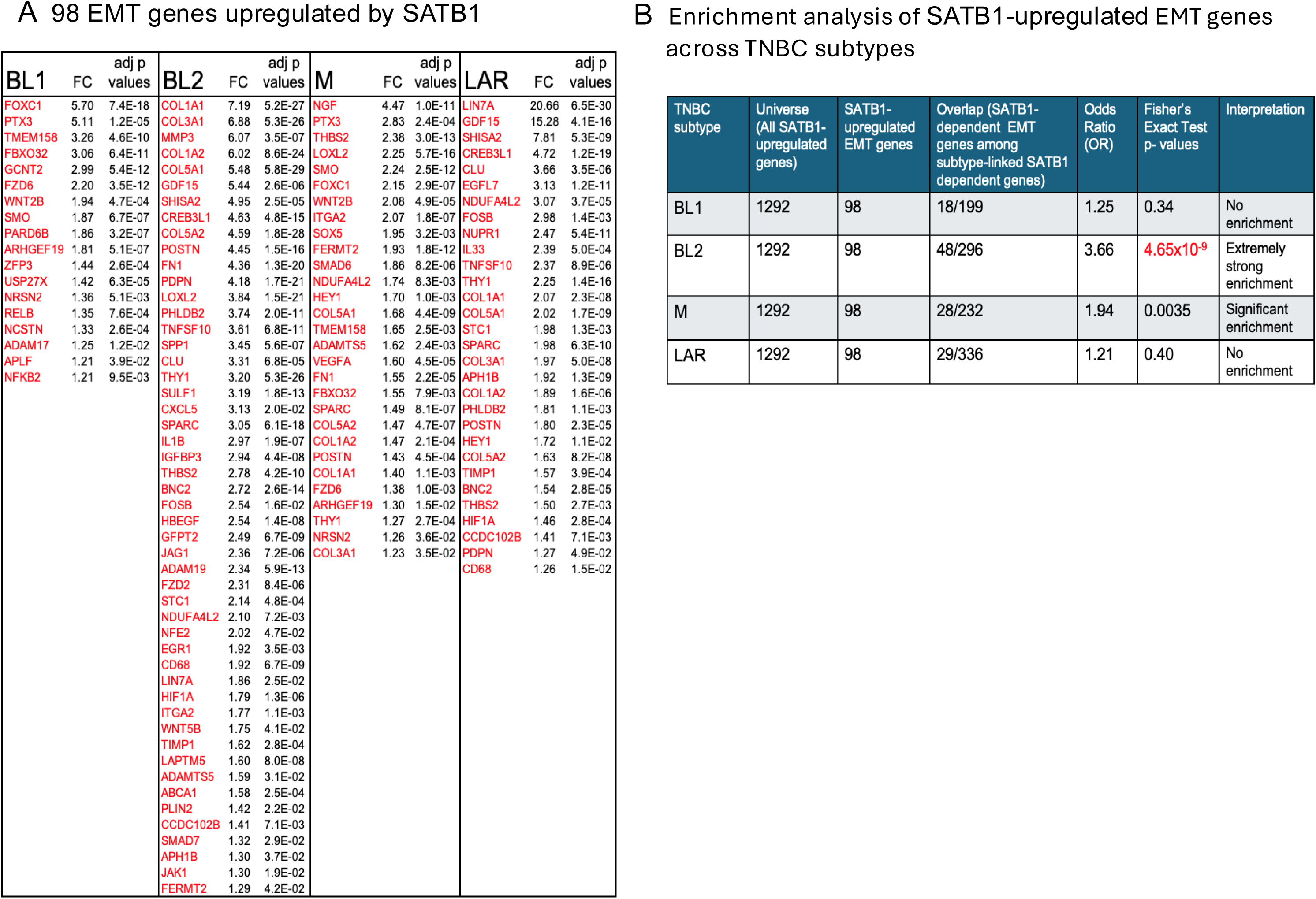
Significant enrichment of 98 SATB1-upregulated EMT genes in the BL2 subtype of TNBC. **A)** Distribution of the 98 EMT genes across the four TNBC subtypes: Basal-like 1 (BL1), Basal-like 2 (BL2), Mesenchymal (M), and Luminal Androgen Receptor (LAR). Fold change (FC) was calculated relative to other TNBC subtypes as described (Lehmann et al., 2021). Adjusted p values (adj P-values) are indicated. B) Enrichment analysis of the 98 SATB1-upregulated EMT genes in each of the four TNBC subtypes using Fisher’s exact test.

To determine whether SATB1-upregulated EMT genes are uniquely enriched in the BL2 subtype, we examined the remaining 202 EMT genes that did not display significant SATB1 dependency by GRO-seq. In contrast to the 98 SATB1-upregulated EMT genes, the 202 SATB1-independent EMT genes showed no enrichment in the BL2 subtype (Odds Ratio: 1.09, Fisher’s exact test, p = 0.61) (see Materials and Methods). Similarly, the BL1 subtype was not enriched in the 202-gene set (Odds Ratio: 0.86, Fisher’s exact test, p = 0.38), whereas the LAR subtype became significantly depleted (Odds Ratio: 0.51, Fisher’s exact test, p = 1.7 × 10⁻⁴). In contrast, the M subtype remained significantly enriched in the 202 EMT genes, similar to the enrichment observed for the 98 SATB1-upregulated EMT genes (Odds Ratio: 1.94, Fisher’s exact test, p = 1 × 10⁻⁵), consistent with the established enrichment of EMT-related genes in the M subtype. These results demonstrate that the 98 SATB1-upregulated EMT genes are uniquely enriched in the BL2 subtype of TNBC, underscoring the selectivity of the SATB1-driven EMT program. Together, these findings indicate that SATB1-regulated EMT genes are preferentially activated in TNBC subtypes associated with aggressive behavior and poor therapeutic response, suggesting that SATB1 drives a defined EMT program that promotes the metastatic potential of breast cancer.

### Global Concordance of B[a]P- and SATB1-Induced Nascent Transcript Profiles

Next, we asked whether the nascent transcript changes induced by SATB1 expression in MCF10A-1 cells are unique to SATB1 or represent a broader response to breast cancer–associated stimuli. To test this, we examined whether brief exposure to benzo[a]pyrene (B[a]P), a well-characterized chemical carcinogen linked to increased breast cancer risk^41,42^, could induce similar nascent transcriptional profiles. Previous studies show that B[a]P promotes migration and invasion in breast epithelial and breast cancer cells, induces mesenchymal markers (vimentin and N-cadherin), represses E-cadherin, and enhances tumor growth and metastasis in mouse models^43–46^. Consistently, MCF10A cells exposed to B[a]P for ∼72 hours (hrs) exhibit increased motility and morphological changes in culture^44^. Therefore, B[a]P was selected as an alternative etiology to compare its transcriptional effects with those induced by SATB1. GRO-seq captured B[a]P-induced transcriptional changes more comprehensively than RNA-seq. After 3 hrs of B[a]P exposure, GRO-seq identified 782 genes, while RNA-seq identified only 129 genes with fc ≥ 2, p ≤ 0.05 (Supplementary Table 6). After 3 days, GRO-seq captured 10 times more genes than RNA-seq, with 1,099 genes compared to 108 genes. This supports the effectiveness of GRO-seq in capturing genes activated by external stimuli.

GSEA revealed strong directional concordance between B[a]P-induced transcriptional changes and SATB1-dependent transcription. Both the top 2,000 B[a]P-upregulated genes and the bottom 2,000 B[a]P-downregulated genes, after either 3 hrs (Figure 4A, B) or 3 days (Figure 4C, D) of B[a]P exposure, were significantly enriched in the SATB1 pre-ranked GRO-seq dataset in the same respective directions (p=0.0001, FDR=0.00015 for both). These results indicate that B[a]P-induced gene expression changes closely mirror the transcriptional program associated with SATB1. Notably, this similarity was unexpected given the brief B[a]P exposure compared with the equilibrated transcriptional state of SATB1-expressing MCF10A-1 cells.

**Figure 4.**
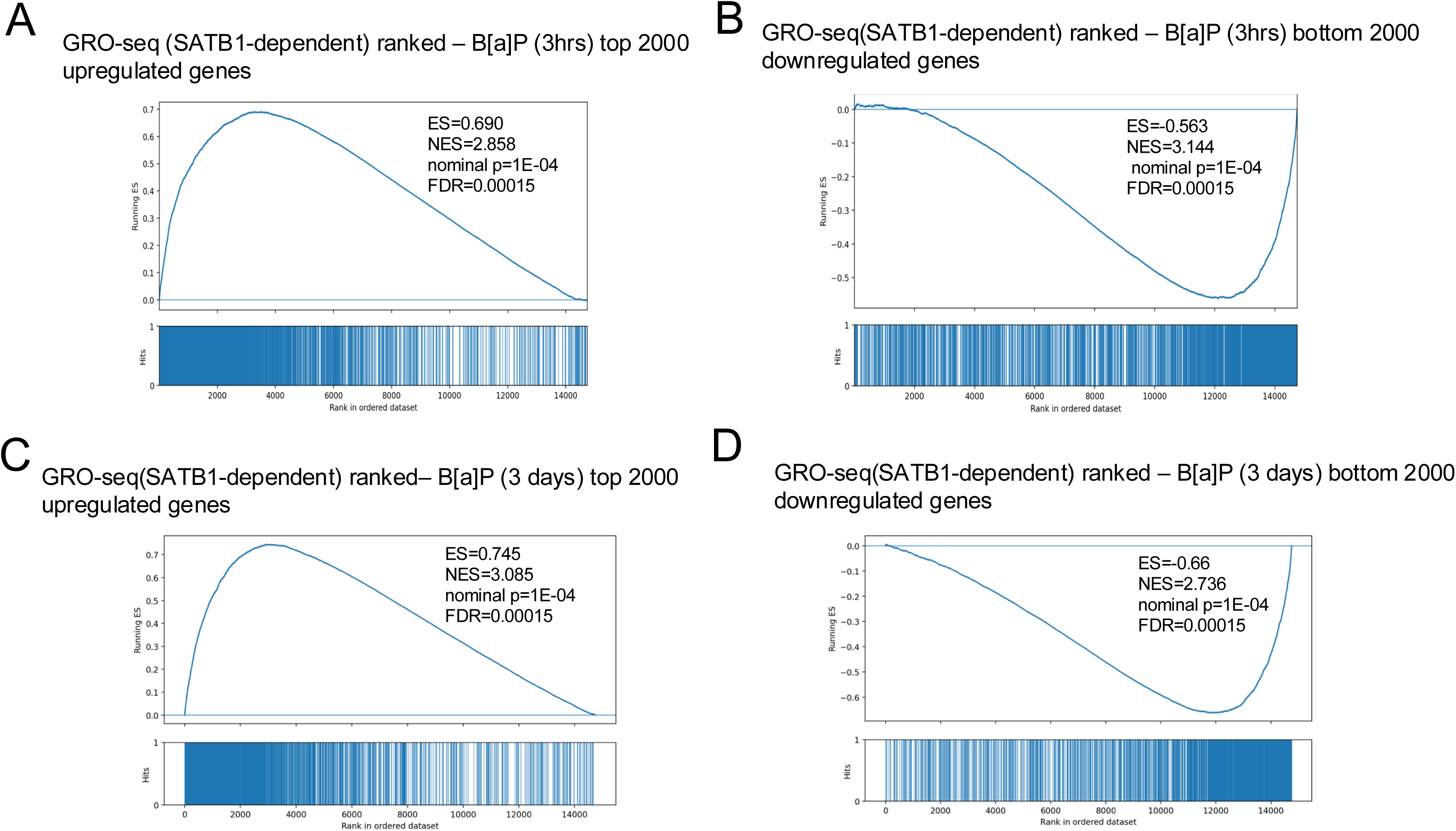
Concordance between SATB1-dependent nascent transcription and B[a]P-induced transcriptional regulation. GSEA was performed using SATB1-dependent nascent transcripts identified by GRO-seq and ranked by log2 fold change. Tested gene sets were defined by either the top 2,000 B[a]P-induced upregulated genes or the top 2,000 downregulated genes. Genes upregulated following B[a]P exposure for 3 hrs (**A**) or 3 days (**B**) were enriched among SATB1-dependent GRO-seq-upregulated genes. Genes downregulated following B[a]P treatment for 3 hrs (**C**) or 3 days (**D**) were enriched among SATB1-dependent GRO-seq-downregulated genes. These results demonstrate statistically significant directional concordance between SATB1-dependent nascent transcriptional regulation and B[a]P-induced changes in nascent transcription.

### B[a]P- and SATB1 upregulate a similar set of EMT genes

Before examining EMT genes among the B[a]P-induced transcription profile, we confirmed whether MCF10A-1 cells properly responded to the B[a]P treatment after 3 hrs. Indeed, B[a]P robustly activated the aryl hydrocarbon receptor (AhR) pathway (Wiki pathway enrichment, p=5.83×10⁻⁶)^118^ as evidenced by the activation of key genes primarily regulated by this pathway, as detected by GRO-seq. These include strong induction of canonical target genes such as *CYP1A1* (fc=163.3, p=5.6×10⁻¹⁰), *AHRR* (fc=33.9, p=5.6 ×10⁻¹¹), *CYP1B1* (fc=20.7, p=2.0×10⁻⁹) and *ARNT* (fc=1.46, p=0.0028). Both *CYP1A1* and *CYP1B1* genes (cytochrome P450 enzymes) are involved in the metabolism and detoxification of xenobiotics like such as B[a]P, often converting it to a carcinogenic metabolite ^119^. The fold change (fc) represents the expression levels in B[a]P-exposed cells relative to non-treated cells.

From a curated set of 300 EMT-promoting genes, B[a]P exposure upregulated 54 EMT genes at 3 hrs and 72 EMT genes at 3 days, using the same thresholds applied to SATB1-expressing cells (fc ≥ 1.2, p ≤ 0.07). Among 72 B[a]P-upregulated EMT genes, 36 coincided with those upregulated by SATB1 (Table 3), showing significant enrichment (Odds ratio: 2.68, Fisher’s exact test: p=4.99 × 10⁻⁴). Gene ontology and pathway analyses revealed that EMT genes induced by B[a]P at 3 days (Table 4) were significantly enriched in functional categories closely mirroring those activated by SATB1 (Table 2), including pathways in cancer, breast cancer, with Benjamini-Hochberg=9.3x10^-11^ or 9.00x 10^-11^, respectively, as well as in other pathways such as cell surface receptor signaling, angiogenesis, transcriptional regulation, and cell migration (Table 4; Figure 5A; genes listed in Supplementary Table 5). Over time (3 hrs to 3 days of B[a]P exposure), B[a]P-dependent changes in the transcription profile evolved by exhibiting many EMT genes with: a) a greater enrichment in functional groups (e.g., cancer pathway, breast cancer, regulation of cell migration, cell surface receptor signaling pathway) (Figure 5B); b) an increased number of activated EMT genes (54 to 72 genes); and c) a greater magnitude of activation (Supplementary Table 7). Representative genes whose expression was significantly increased during this period (3 hrs to 3 days) include *WNT4, FBX032, TNFSF10, ABCA1, FGF1, BMP7, GJA1, HBEGF, WNT10A, GDF15, NFE2, ELF3, EPHA4, ERG, CEMIP, WNT9B, CCN4, PDGFD,* and *JAG1* (Table 3).

**Figure 5.**
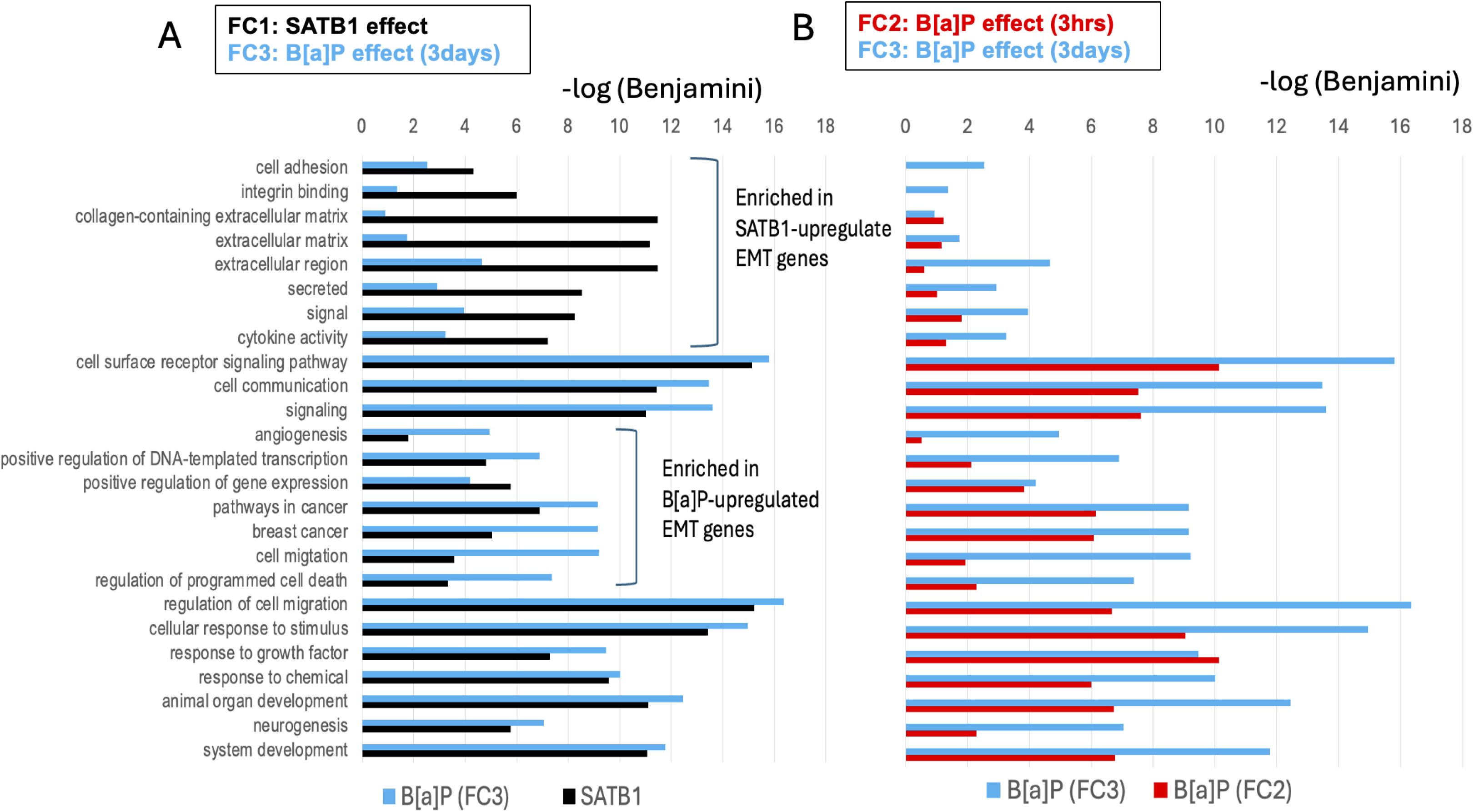
DAVID gene enrichment analysis of B[a]P-upregulated genes compared with SATB1-upregulated genes identified by GRO-seq in SATB1-stably transduced MCF10A-1 cells. **A)** Gene enrichment within each DAVID functional group is shown for B[a]P-upregulated EMT-promoting genes after 3 days of treatment (72 genes, FC3; blue bars) and SATB1-upregulated EMT-promoting genes (98 genes, FC1; black bars). **B)** Gene enrichment is shown for B[a]P-upregulated EMT-promoting genes after 3 days of treatment (72 genes, FC3; blue bars) and after 3 hrs of treatment (54 genes; red bars), demonstrating a time-dependent increase in enrichment of B[a]P-upregulated EMT-promoting genes within each gene group. Whereas SATB1-upregulated genes are highly enriched in cellular components related to extracellular matrix (ECM) structure and function (top six gene groups), these gene groups remain weakly enriched among B[a]P-upregulated genes after 3 days of B[a]P exposure. The x-axis [−log(Benjamini)] indicates the statistical significance of each gene group.

**Table 3.** 72 B[a]P-upregulated EMT-promoting genes from GRO-seq.

| Gene ID | fold change | Gene ID | fold change | Gene ID | fold change | Gene ID | fold change |
| --- | --- | --- | --- | --- | --- | --- | --- |
| **WNT4** | 39.15 | **VANGL2** | 4.97 | **PHLDB2** | 2.68 | **GAB1** | 1.67 |
| **FBXO32** | 35.29 | **ELF3** | 4.70 | **AGER** | 2.34 | **TCF4** | 1.65 |
| **TNFSF10** | 22.47 | **FZD10** | 4.49 | **NFKB2** | 2.32 | **IL1B** | 1.65 |
| **ABCA1** | 12.34 | **EPHA4** | 4.26 | **PARD6B** | 2.16 | **SDCBP** | 1.63 |
| **SOX5** | 12.34 | **GFPT2** | 4.15 | **ID2** | 2.12 | **FGFR2** | 1.63 |
| **FGF1** | 11.27 | **ERG** | 4.11 | **RGCC** | 2.07 | **NFKB1** | 1.59 |
| **FOSB** | 10.39 | **GSDMB** | 4.04 | **RAP2B** | 2.03 | **NCSTN** | 1.58 |
| **MAP2K6** | 8.63 | **CEMIP** | 3.97 | **EFNA1** | 2.02 | **FGFR1** | 1.52 |
| **BMP7** | 8.14 | **PIK3R3** | 3.94 | **PIK3CA** | 2.01 | **SMAD7** | 1.49 |
| **GJA1** | 7.79 | **WNT9B** | 3.77 | **FGFR3** | 1.98 | **SLC39A6** | 1.46 |
| **HBEGF** | 7.00 | **COL5A2** | 3.55 | **APH1B** | 1.97 | **ACVR1** | 1.43 |
| **WNT10A** | 6.99 | **PIK3C2G** | 3.54 | **PDGFD** | 1.89 | **SLC39A10** | 1.42 |
| **GDF15** | 6.94 | **CCN4** | 3.45 | **NOTCH1** | 1.89 | **KLF8** | 1.39 |
| **IL13** | 6.51 | **APLF** | 3.44 | **ITGA2** | 1.83 | **ADAM17** | 1.39 |
| **SULF1** | 6.44 | **PIK3C2B** | 2.98 | **HIF1A** | 1.81 | **TCF7L2** | 1.38 |
| **EGR1** | 6.22 | **REL** | 2.82 | **JAK1** | 1.75 | **MET** | 1.33 |
| **NUPR1** | 5.66 | **FZD6** | 2.75 | **TWIST1** | 1.73 | **MAP2K5** | 1.27 |
| **NFE2** | 5.06 | **USP27X** | 2.68 | **JAG1** | 1.67 | **SOS2** | 1.24 |

**Table 4.** B[a]P-upregulated EMT-promoting genes (GRO-seq)

| Pathways in cancer | Breast cancer | collagen-containing extracellular matrix | extracellular region | cell adhesion | Integrin binding | angio-genesis | positive regulation of DNA-templated transcription | regulation of cell migration<br>↔ |  | regulation of programmed cell death | signal transduction | response to growth factor | animal organ development<br>↔ |  |
| --- | --- | --- | --- | --- | --- | --- | --- | --- | --- | --- | --- | --- | --- | --- |
| FGF1 | FGF1 | BMP7 | AGER | ADAM17 | ADAM17 | EFNA1 | ACVR1 | ACVR1 | RGCC | ADAM17 | CCN4 | ACVR1 | ADAM17 | NCSTN |
| FGFR1 | FGFR1 | COL5A2 | BMP7 | ACVR1 | CCN4 | FGF1 | BMP7 | AGER | SDCBP | ACVR1 | ERG | BMP7 | ACVR1 | NFKB2 |
| FGFR2 | FZD10 | FGFR2 | CCN4 | AGER | FGF1 | FGFR1 | EGR1 | BMP7 | SMAD7 | BMP7 | FGF1 | EGR1 | AGER | NOTCH1 |
| FGFR3 | FZD6 | GDF15 | CEMIP | BMP7 | IL1B | FGFR2 | ELF3 | CCN4 | SULF1 | EFNA1 | GAB1 | FGF1 | BMP7 | NUPR1 |
| FZD10 | JAG1 | ITGA2 | COL5A2 | CCN4 | ITGA2 | GAB1 | ERG | CEMIP | TWIST1 | EGR1 | GDF15 | FGFR1 | CCN4 | PARD6B |
| FZD6 | NFKB2 | SULF1 | EFNA1 | EFNA1 |  | HIF1A | FGF1 | EFNA1 | WNT4 | FGFR1 | GJA1 | FGFR2 | COL5A2 | PIK3CA |
| HIF1A | NOTCH1 |  | FGF1 | EPHA4 |  | JAG1 | FGFR2 | EPHA4 |  | FGFR2 | HBEGF | FGFR3 | EFNA1 | PIK3R3 |
| IL13 | PIK3CA |  | FGFR1 | FGFR2 |  | NOTCH1 | FOSB | FGF1 |  | HIF1A | HIF1A | GAB1 | EGR1 | SDCBP |
| ITGA2 | PIK3R3 |  | FGFR2 | IL1B |  | PIK3CA | HIF1A | FGFR1 |  | IL13 | IL13 | GDF15 | EPHA4 | SMAD7 |
| JAG1 | SOS2 |  | FGFR3 | ITGA2 |  |  | ID2 | GAB1 |  | IL1B | IL1B | MAP2K5 | FGF1 | SOX5 |
| JAK1 | TCF7L2 |  | GDF15 | JAG1 |  |  | IL1B | GDF15 |  | MAP2K5 | MAP2K5 | NOTCH1 | FGFR1 | SULF1 |
| MET | WNT10A |  | HBEGF | NOTCH1 |  |  | JAG1 | GJA1 |  | MAP2K6 | MAP2K6 | PDGFD | FGFR2 | TCF4 |
| NFKB1 | WNT4 |  | IL13 | RAP2B |  |  | MAP2K5 | HBEGF |  | MET | MET | PIK3CA | FGFR3 | TCF7L2 |
| NFKB2 | WNT9B |  | IL1B |  |  |  | MET | HIF1A |  | NFKB1 | NFKB1 | SMAD7 | FZD10 | TNFSF10 |
| NOTCH1 |  |  | JAG1 |  |  |  | NFE2 | IL1B |  | NOTCH1 | RAP2B | SOX5 | FZD6 | TWIST1 |
| PIK3CA |  |  | MET |  |  |  | NFKB1 | ITGA2 |  | NUPR1 | TNFSF10 | TWIST1 | GAB1 | VANGL2 |
| PIK3R3 |  |  | NFKB1 |  |  |  | NFKB2 | JAG1 |  | PIK3CA |  | WNT10A | GJA1 | WNT10A |
| SOS2 |  |  | NOTCH1 |  |  |  | NOTCH1 | MAP2K5 |  | PIK3R3 |  | WNT4 | HIF1A | WNT4 |
| TCF7L2 |  |  | PDGFD |  |  |  | NUPR1 | MET |  | RGCC |  |  | ID2 | WNT9B |
| WNT10A |  |  | SDCBP |  |  |  | REL | NOTCH1 |  | SLC39A10 |  |  | IL1B |  |
| WNT4 |  |  | TNFSF10 |  |  |  | RGCC | PARD6B |  | TCF7L2 |  |  | ITGA2 |  |
| WNT9B |  |  | WNT10A |  |  |  | TCF4 | PDGFD |  | TNFSF10 |  |  | JAG1 |  |
|  |  |  | WNT4 |  |  |  | TCF7L2 | PHLDB2 |  | TWIST1 |  |  | MAP2K5 |  |
|  |  |  | WNT9B |  |  |  | TWIST1 | PIK3R3 |  | USP27X |  |  | MAP2K6 |  |
|  |  |  |  |  |  |  | WNT4 | RAP2B |  | WNT4 |  |  | MET |  |

Important genes commonly upregulated by B[a]P and SATB1 include *FBX032, ABCA1, SOX5. HBEGF, GDF15, NFE2, CEMIP, CCN4, ITGA2, HIF1A, JAG1, and MET*. On the other hand, SATB1 uniquely activated genes are closely linked to extracellular matrix remodeling and the development of metastatic competence, such as *MMP9*, *VEGFA*, *FN1*, *FOXC1*, *HEY1*, SMO and CXCR5. Therefore, at least on day 3 of B[a]P treatment, a notable difference is that B[a]P-induced EMT genes showed weak enrichment for ECM-related functional categories, including collagen-containing extracellular matrix, integrin binding, cytokine activity, and proteolytic ECM remodeling (Figure 5A). Proteases critical for ECM degradation (MMPs, ADAMs, ADAMTS) were also absent from the B[a]P-induced gene set. While a longer exposure time could yield valuable information on ECM-related genes, we opted not to extend the cell culture for these experiments, as we observed that control MCF10A-1 cells tend to shift toward malignant-like phenotypes with prolonged culture^2^.

DAVID functional annotation and pathway analyses of B[a]P-upregulated and SATB1-upregulated genes identified key angiogenic regulators, such as *JAG1*^120^ and *HIF1A*^121^ (Table 2 and 4), as well as *MET*^95,96^, were commonly upregulated. In breast cancer, hypoxia upregulates JAG1 through activation of HIF1A and HIF1A directly activates MET transcription, and these events further promote tumor angiogenesis, immune evasion, and resistance to therapy^122^. Notably, B[a]P preferentially activated components of the FGF/FGFR signaling pathway (*FGF1*, *FGFR1*, *FGFR2*), which also plays crucial roles in angiogenesis (Table 4, Supplementary Table 5). Dysregulation of this pathway can lead to resistance to cancer treatments^123^. These results indicate that even brief exposure (e.g. 3 days) to B[a]P activates key regulators of angiogenesis commonly found in the SATB1-regulated EMT programs and pathways for angiogenesis.

With regards to downregulated EMT genes, B[a]P significantly downregulated *DKK1, EMP2, CDH1, and OVOL2*, similar to SATB1 effects, among the 27 EMT-inhibiting gene list. Additionally, the GRO-seq analysis revealed that among the metastasis suppressor genes, *RHOGD12 (ARHGDIB)* is significantly downregulated by B[a]P treatment, both at 3 hrs (fc=0.42, p=0.00096) and on day 3 (fc=0.39, p=0.0006). Similarly, CLDN4 (*Claudin-4*) is downregulated by B[a]P at 3 hrs (fc=0.51, p=0.03) and on day 3 (fc=0.39, p=0.007). These genes were also repressed in SATB1-expressing cells. Furthermore, B[a]P significantly downregulated *SSeCKS* (or *AKAP12*) with values of (fc=0.09, p=0.005 at 3 hrs) and (fc=0.059, p=0.004 on day 3). Additionally, *DLC1* and *DRG1* were downregulated solely by B[a]P: *DLC1* at 3 hrs (fc=0.40, p=0.0001) and *DRG1* at 3 hrs (fc=0.72, p=0.03) and on day 3 (fc=0.74, p=0.05).

Together, these results indicate that exposure to an environmental carcinogen and SATB1 expression activate overlapping transcriptional trajectories associated with EMT. SATB1 engages a comprehensive EMT program that prominently features extracellular matrix remodeling, while B[a]P induces a temporally progressive EMT response with comparatively modest transcriptional effects on ECM remodeling, which expand in scope and magnitude with prolonged exposure. This convergence supports the existence of a shared metastatic gene framework that can be initiated by distinct oncogenic insults.

### SATB1-expression diminished B[a]P Effects

Finally, we asked whether B[a]P treatment further enhances SATB1-driven EMT gene expression or whether B[a]P-induced effects are attenuated in the presence of SATB1. To address this, we compared the transcriptional responses of EMT genes to B[a]P exposure in MCF10A-1 cells with or without SATB1 transduction at two time points: 3 hrs [B[a]P effects in SATB1-expressing cells (FC4) versus B[a]P alone (FC2)] and 3 days [B[a]P effects in SATB1-expressing cells (FC5) versus B[a]P alone (FC3)].

After 3 hrs of B[a]P treatment SATB1-expressing cells further increased nascent transcription of multiple EMT genes, including *COL1A1*, *COL1A2*, *COL3A1*, *IGFBP3*, *SPARC*, *GJA1*, *PRRX1*, and *BMP4* (FC4), many of which were already strongly induced by SATB1 alone (Figure 6A). In contrast, this enhancement was largely diminished after 3 days of B[a]P exposure, with the notable exception of *IL6* (Figure 6B). This pattern differed markedly from control MCF10A-1 cells, in which B[a]P exposure produced a pronounced increase in EMT gene activation at day 3 (FC3), compared with the more modest effects observed in SATB1-expressing cells at the same time point (FC5) (Figure 6C).

**Figure 6.**
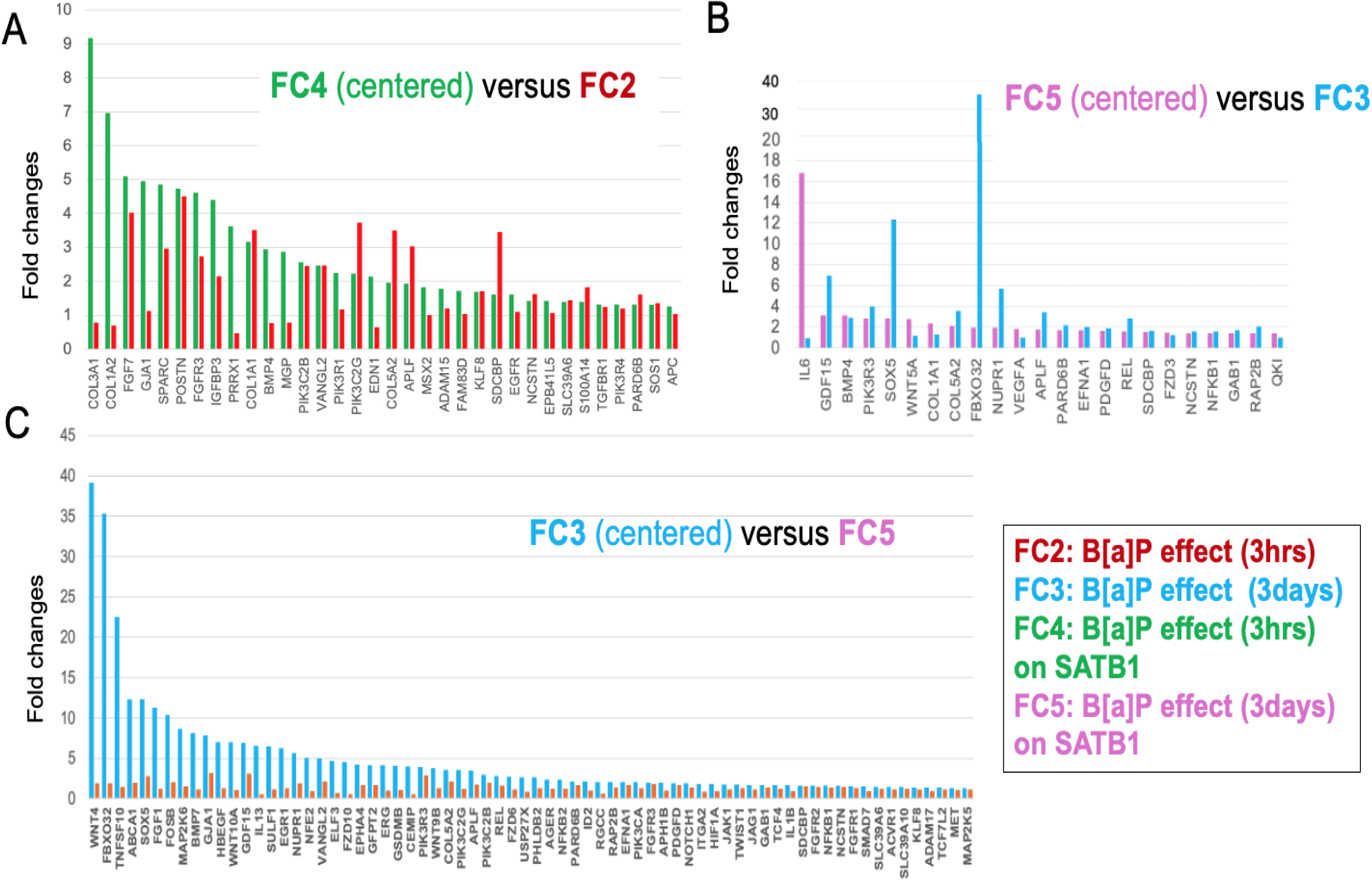
Time-dependent reduction in the effects of B[a]P on EMT gene activation in the presence of SATB1, as detected by GRO-seq. **A)** Genes upregulated by B[a]P after 3 hrs in SATB1-expressing MCF10A-1 cells (selected using fc > 1.2 and p < 0.07, and arranged in descending order centered on FC4; green bars) were compared with genes upregulated by B[a]P after 3 hrs in control MCF10A-1 cells (without selection; FC2, red bars). Fold changes in transcript levels are shown for each gene. **B)** Genes upregulated by B[a]P after 3 days in SATB1-expressing MCF10A-1 cells (selected using fc > 1.2 and p < 0.07, and arranged in descending order centered on FC5; pink bars) were compared with genes upregulated by B[a]P after 3 days in control MCF10A-1 cells (without selection; FC3, blue bars). **C)** Genes upregulated by B[a]P after 3 days in control MCF10A-1 cells (selected using fc > 1.2 and p < 0.07, and arranged in descending order centered on FC3; blue bars) were compared with genes upregulated by B[a]P after 3 days in SATB1-expressing MCF10A-1 cells (without selection; FC5, red bars). This analysis is similar to panel B, except that transcript levels for FC3 genes were centered instead of FC5 genes. Fold changes in transcript levels are shown for each gene.

Consistent with this observation, both the number and magnitude of EMT genes affected by B[a]P were reduced in SATB1-expressing cells at day 3. Only 26 EMT genes were significantly altered in the FC5 condition (fc ≥ 1.2, and p ≤ 0.07), compared with 72 EMT genes in the FC3 condition, and activation levels were generally modest (<3-fold), except for *IL6*, which was induced ∼17-fold (p = 3 × 10⁻⁶). Notably, *IL6* represents a unique case in which neither SATB1 nor B[a]P alone induced expression, but robust activation occurred upon B[a]P treatment in a SATB1-expressing context.

*IL6* encodes a pro-inflammatory cytokine known to promote EMT and metastatic potential in breast cancer through autocrine and paracrine signaling mechanisms^124,125^. Tumor-derived IL6 contributes to a pro-tumorigenic microenvironment that facilitates invasion and metastasis, including dissemination to bone and lung. Consistent with prior reports, B[a]P-induced activation of *IL6* has been linked to inflammatory microenvironments that enhance migration and metastatic behavior of breast epithelial cells^45^.

Together, these analyses indicate that while B[a]P can temporarily enhance EMT gene transcription in SATB1-expressing cells, its effects largely diminish with prolonged exposure. This attenuation likely reflects SATB1’s dominant role in establishing and maintaining the EMT-associated transcriptional program, into which B[a]P-induced signals are subsequently integrated. Thus, once SATB1-driven EMT programs are established, further exposure to carcinogens contributes minimally to additional EMT gene activation, with the notable exception of inflammatory signaling pathways. Therefore, after SATB1-driven EMT transcriptional programs are in place, B[a]P exposure has only a limited impact on EMT genes, indicating that SATB1 functions as a primary organizer of EMT gene expression in this context.

## Discussion

Hundreds of genes are transcriptionally reprogrammed during breast cancer progression toward metastasis. SATB1 is a central driver of this reprogramming when aberrantly expressed in cancer cells; however, the key downstream gene networks through which SATB1 promotes metastasis have not been systematically defined. In this study, we addressed this gap by focusing on EMT, a multifaceted process that promotes invasion, dissemination, metastatic competence, immune escape and therapeutic resistance.

Among a curated set of 300 EMT-promoting genes, we identified 98 that were significantly upregulated by SATB1 in breast epithelial cells exhibiting SATB1-induced malignancy. These findings suggest that malignant progression is driven by a defined EMT transcriptional program orchestrated by SATB1. This conclusion is further supported by the significant overlap between SATB1-regulated EMT genes and those induced by carcinogen exposure, indicating that distinct oncogenic etiologies largely converge on a common EMT-promoting program rather than activating random subsets of EMT genes. Importantly, these 98 SATB1-activated EMT genes were selectively enriched in the aggressive BL2 subtype of TNBC, recognized as one of the most aggressive TNBC subtypes, underscoring their potential clinical relevance.

### Concordant nascent transcripts and EMT genes upregulated by SATB1 and B[a]P

B[a]P, a prototypical environmental carcinogen, induces malignant transformation in non-malignant breast epithelial cells in vitro and promotes the metastasis of breast cancer cells in vivo in xenograft models^43,44^. GSEA revealed a high concordance of genes regulated by SATB1 and B[a]P, both in upregulation and downregulation (Figure 4). At the individual gene level, B[a]P exposure activated 72 of the 300 EMT-promoting genes within three days, 36 of which overlapped with SATB1-upregulated EMT genes. This represents a significant enrichment of EMT genes upregulated by SATB1 among those upregulated by B[a]P (Fisher’s exact test, p ≈ 10⁻^4^). Functional annotation and pathway analyses revealed striking similarities between EMT genes activated by SATB1 and those induced by B[a]P (see Table 2 vs. Table 4 and Supplementary Table 5). The only exception was a weaker enrichment of extracellular matrix genes (i.e., genes relevant to ECM structure and function) following B[a]P exposure, which may become more pronounced with longer treatment. This substantial overlap is particularly noteworthy given the distinct etiologies of tumor development: direct oncogenic reprogramming by the chromatin organizer SATB1 through stable expression in cultured cells in one context, versus cancer initiation triggered by brief exposure to a chemical carcinogen in the other.

Over time, from 3 hrs to 3 days, B[a]P-induced EMT gene activation intensified and expanded, increasing its overlap with the 98 genes upregulated by SATB1. This supports the notion that the SATB1-upregulated EMT genes form a core EMT program associated with malignant progression.

Importantly, B[a]P did not act synergistically with SATB1; rather, its transcriptional effects were significantly diminished in SATB1-expressing cells. This suggests that once established, the SATB1-driven transcriptional program predominates and integrates various oncogenic inputs into a cohesive EMT network. Together, these findings indicate that the 98 EMT genes represent a central, regulator-defined transcriptional program underlying metastatic competence in breast cancer cells.

### Potential mechanisms for EMT gene activation in SATB1-expressing MCF10A-1 cells

To understand how a large number of EMT genes may be regulated in SATB1-expressing MCF10A-1 cells, we can draw insights from our recent findings on how SATB1 organizes the genome to regulate hundreds of genes in a cell type- or cell function-specific manner. We recently employed a modified ChIP-seq approach using urea-purified crosslinked chromatin to detect direct SATB1 binding sites^12^. In contrast, conventional ChIP-seq primarily detects indirect SATB1 binding sites. Based on these results, we proposed that SATB1 establishes a novel “two-tiered” chromatin organization^12^. This organization consists of (1) an insoluble, extraction-resistant “subnuclear architectural structure” tier, where SATB1 binds directly to BURs in the genome to form a stable chromatin scaffold, and (2) a soluble, accessible chromatin tier corresponding to gene-rich regions, including enhancers, with which SATB1 interacts indirectly, presumably through associations with other factors bound to accessible chromatin regions. These two tiers are interconnected, as SATB1-bound BURs in the subnuclear structure extensively interact with accessible chromatin encompassing multiple gene-rich regions. In thymocytes, these BUR-gene-rich region interactions are linked to gene expression^12^, as SATB1 deletion abolished both the interactions and proper gene expression. SATB1 is known to associate with transcription factors and chromatin-modifying proteins^1,3,11,14,23,126,127^, and this property likely facilitates the assembly of transcriptional complexes at SATB1 target loci on the SATB1 chromatin scaffold to regulate gene expression. Although the precise mechanisms remain unresolved, SATB1-dependent activation of EMT genes in MCF10A-1 cells is likely mediated through this two-tiered chromatin architecture, whereby SATB1-bound BURs spatially coordinate distantly located EMT gene loci into shared transcriptional hubs.

Even in the absence of SATB1, short-term exposure to B[a]P upregulated 36 of the 98 EMT genes normally activated by SATB1. This effect is attributable to B[a]P itself rather than to SATB1 induction during the short experimental window, as SATB1 mRNA induction was not detected by either RNA-seq or GRO-seq analyses. B[a]P rapidly activated the AhR signaling pathway in MCF10A-1 cells, as indicated by Reactome analysis (p = 0.0055 for GRO-seq data), inducing cytochrome P450 genes such as CYP1A1 and CYP1B1. Upon activation, AHR translocates to the nucleus, where it can induce EMT gene expression in a context-dependent manner^128^. Because CYP1A1, CYP1B1, ARNT2, and AHRR are also upregulated in a SATB1-dependent manner, SATB1 likely engages the AhR signaling pathway, thereby contributing to the shared induction of EMT genes with B[a]P.

### Increased Variability of Nascent Transcription in SATB1-Expressing

One plausible explanation for the increased variability in nascent transcription observed in SATB1-expressing cells is that SATB1 alters the dynamics of transcriptional bursting. Transcription is a complex and highly regulated process involving multiple dynamic steps, such as the recruitment of RNA Polymerase II (Pol II) to gene loci, its residence time, and elongation^129,130^. Genes are transcribed in discrete bursts of activity, alternating between short periods of active transcription and longer periods of inactivity—a phenomenon known as transcriptional bursting. This bursting contributes to intrinsic variability in nascent transcripts (transcriptional noise) across species^131,132^, resulting in significant cell-to-cell heterogeneity in Pol II residence time along genes within a cell population^133^. Transcriptional burst frequency and amplitude of transcriptional bursts are influenced by chromatin accessibility, interactions with regulatory factors, and higher-order genome organization^134^.

SATB1 actively participates in chromatin organization by tethering genomic elements (BURs) to multiple gene loci. Since dynamic genome interactions have been observed at the single-cell level in super-resolution imaging studies^135–137^, SATB1-mediated chromatin interactions are likely to be highly dynamic. Through this three-dimensional chromatin organization, SATB1 may enhance locus-specific variability in Pol II engagement across cells, thereby amplifying cell-to-cell differences in nascent RNA levels detected by GRO-seq. The hypothesis that SATB1 influences transcriptional bursting can be tested in future studies. Importantly, intrinsic transcriptional variability can be biologically advantageous and has been linked to processes such as development and environmental adaptation^138^ ^139^. The increased variability in Pol II activity in SATB1-expressing cells may therefore reflect a chromatin environment poised for change or undergoing active transcriptional reprogramming. This feature may be critical for SATB1’s function in altering cell states, such as during development and cancer progression, both of which require substantial shifts in gene expression programs.

Despite the increased variability in nascent transcription, multiple independent lines of evidence support the existence of a coherent SATB1-dependent EMT program identified by GRO-seq. These include: (a) strong concordance between the 300 EMT genes and genes upregulated in both GRO-seq and RNA-seq datasets (p = 0.0001 for each); (b) similar pathway enrichment and strong functional overlap between SATB1- and B[a]P-induced EMT gene sets identified by GRO-seq; (c) significant overlap of EMT genes across distinct oncogenic contexts; and (d) selective enrichment of the 98 SATB1-upregulated EMT genes in a TNBC subtype associated with the highest risk for disease progression and with the shortest median survival of 2.4 years^140^. Collectively, these findings suggest that the observed transcriptional variability reflects an intrinsic feature of SATB1-mediated chromatin reorganization rather than a limitation in identifying a biologically meaningful EMT program.

### Future perspectives: SATB1-target EMT gene signature as prognostic tools

High SATB1 protein expression is strongly correlated with poor prognosis across various cancer types, including breast cancer. In contrast, total *SATB1* transcript levels exhibit variable expression in normal breast tissue. As a result, SATB1-regulated EMT genes, which are downstream targets of SATB1, may offer a biologically relevant gene signature for assessing metastatic risk. Preliminary analysis of RNA-seq data from 32 TNBC patient-derived xenografts^141^ revealed that metastatic tumors do not uniformly overexpress a small subset of top SATB1-induced EMT genes. Instead, distinct combinations of EMT genes are activated in individual tumors, suggesting that metastatic potential is determined by the breadth and intensity of EMT program activation rather than a few dominant markers (unpublished results). Therefore, screening only a limited subset of EMT genes is unlikely to provide reliable prognostic power. A larger EMT gene panel is better suited to capture tumor heterogeneity and metastatic risk.

Current breast cancer genomic assays primarily stratify recurrence risk based on tumor proliferation and hormone responsiveness, with relatively few genes directly linked to metastatic competence. Notably, assays such as MammaPrint^142,143^ and Oncotype DX^144,145^ were developed and validated primarily for ER-positive, HER2-negative breast cancer and predominantly include genes related to proliferation. In contrast, the SATB1-activated EMT gene program defines a distinct transcriptional axis of disease biology, capturing metastatic competence through the coordinated regulation of EMT-related processes, including invasion, extracellular matrix remodeling, cellular plasticity, drug resistance, and immune evasion. Currently, there is no widely adopted or clinically validated gene signature that reliably stratifies TNBC patients into distinct low-risk and high-risk outcome groups^114^. Furthermore, TNBC molecular subtyping is not a standard-of-care biomarker and does not directly guide treatment decisions in most clinical settings. There is a need for biologically grounded prognostic tools that reliably reflect metastatic potential.

Approximately 1,000 coding and non-coding transcripts related to EMT or MET (mesenchymal-to-epithelial transition) have been curated from the literature, including 810 protein-coding genes in the EMTome dataset^146^ and 1,101 protein-coding genes in the dbEMT 2.0 database^147^. These databases serve as valuable reference resources for analyzing EMT-associated signatures, with some EMT genes linked to poor outcomes in certain cancers^146,148^. Among the 98 SATB1-induced EMT genes, 44 overlap with EMTome and 38 with dbEMT 2.0, sharing 25 genes between the two databases. A key distinction between our EMT gene set and these collections is that our genes are regulator-defined, directly activated by SATB1, rather than a compilation of EMT-associated genes from diverse experimental contexts. Given that SATB1 is frequently detected in metastasis-capable tumors across multiple human cancers, EMT-promoting genes directly activated by SATB1 are likely to constitute an effector transcriptional program critical for metastatic progression. We anticipate that this SATB1-dependent program is preferentially activated in TNBC tumors primed for metastasis and, after full validation in large clinical cohorts, may serve as a predictive tool for metastatic risk in TNBC and potentially other breast cancer subtypes.

## Materials and Methods

### Cell lines

We used the MCF10A-1 cell line described in Ordinario et al^2^.This cell line has been derived from MCF10A cells, originally obtained from the American Type Culture Collection (ATCC), by passaging more than 50 times in culture medium containing fetal bovine serum. MCF10A-1 cells stably transfected with either the pLXSN retroviral vector (control cells) or SATB1 the pLXSN-SATB1 construct (SATB1 expressing cells) were used and maintained as described ^2^.

### Benzo[a]Pyrene Treatment

MCF10A-1 (control) and SATB1-expressing MCF10A-1 cells were treated with 4uM benzo[a]pyrene (Sigma) for either 3 hrs or 3 days, referring to Genies et al^149^. After treatment, cells were washed three times with PBS.

### RNA-seq Library Preparation and Sequencing

Total RNA was purified by TRI reagent (Sigma) and followed by the RNeasy kit (Qiagen). One microgram of total RNA from each sample was reverse transcribed using the QuantiTect Rev Transcription kit (Qiagen). The quality of total RNA was confirmed by 2100 bioanalyzer (Agilent Technologies, Inc.) and high-quality RNA samples (RIN>8) were further processed at QB3-Berkeley Genetics Core Facilities (University of California, Berkeley). After mRNA capture using Oligo(dT)25 magnetic beads (ThermoFisher), the library for sequencing was prepared on Apollo 324TM with PrepXTM RNA-Seq Library Preparation Kits (WaferGen Biosystems, Fremont, CA) according to the manufacturer’s recommendation. RNA sequencing was performed for triplicates per sample on HiSeq4000 machine (illumina, San Diego, CA) generating on average 30 million reads per sample.

### GRO-seq Library Preparation and Sequencing

GRO-seq library construction was prepared by applying the protocol for RNA preparation of Gro-seq (Methods in Molecular Biology, vol. 1468, Chapter 9, 2017, by A. Gardini; all recipes were in it), followed by library construction using Takara SMATer-kit. Briefly, nuclei were isolated from tissue culture cells, counted the number, washed and frozen (1 x 10^6 of nuclei in 10 ul of freezing buffer) at -80C until ready for *in vitro* Nuclear Run-on (NRO) transcription reaction. When the Gro-seq preparation was ready, 100 ul of nuclei stock were mixed with 100 ul of 2x NRO buffer (containing, ATP, GTP, BrUTP and CTP) and incubated at 37C for 7 min. The reaction was stopped by adding TRI-LS reagent (Sigma) and purified RNA in 20 ul of water with RNase inhibitor (1U/ul). The collected RNAs were further treated with DNase I (Turbo DNA-free kit, Bio-Rad), RNA fragmentation by RNA fragmentation kit (Ambion), purified by Micro-Bio-Spin P-30, incubated with T4 polynucletide kinase (PNK) at 37C for 60 min, and inactivated the PNK at 75C for 5 min. The final Br-UTP incorporated RNAs were immunoprecipitated with beads (anti-BrdU antibody conjugated agarose; Santa Cruz; sc-32323 AC), washed beads, and eluted by 100 ul of 0.25M EDTA-TE for 4 times. The eluted RNAs were purified by EtOH precipitation, and the final RNAs were dissolved in 15 ul of water and quantitated by Qubit reading, followed by Library construction. Library construction was performed by Takara SMATer smRNA library kit as manufactured suggested (polyadenylation, cDNA synthesis, PCR and clean up). Library sequencing was performed in UCSF sequence facility using Illumina High-seq 4000.

### Analysis of RNA-seq and GRO-seq

#### RNA-seq data analysis

Raw sequencing data went through quality control steps to remove technical features such as Illumina adapter, polyAT sequences, PhiX contamination, using HTStream (v1.0.0, https://github.com/s4hts/HTStream). This removed technical features such as Illumina adapters, polyAT sequences, PhiX contamination, and reads less than 21 bp in length. Reads were then aligned to GRCh38 reference genome using STAR aligner^150^. Raw counts generated from STAR aligner were normalized using TMM method in edgeR^151^. Differential expression analysis was carried out using limma+voom^152,153^. For differential expression testing, the genomic alignments were restricted to those that map uniquely to the set of known Ensembl IDs (this includes the exons of all protein coding mRNAs as well as many other coding and noncoding RNAs). STAR aggregates this subset of mappings on a per gene basis as raw input for the program limmat+voom. False Discovery Rate was controlled using Benjamini-Hochberg procedure^154^. Genes were filtered using fold changes and p values (fc ≥ 1.2 or fc ≤ 2 as described, and *adj p* ≤ 0.05).

#### GRO-seq data analysis

Raw sequencing data went through quality control steps to remove technical features such as Illumina adapter, polyAT sequences, PhiX contamination, using HTStream (v1.0.0, https://github.com/s4hts/HTStream). Reads less than 21 bp in length were excluded from downstream analysis. Reads were aligned to the GRCh38 human genome using the MEM algorithm in BWA (v0.7.16a)^155^, allowing a counts table to be generated using featureCounts^156^ and the corresponding genome annotation (GENCODE v34). High quality mapping that has a quality score greater than 10 was used for downstream analysis. (Note: The gene expression was also quantified using groHMM^157^, with the first 200 bp downstream of the transcription start site (TSS) excluded to account for RNA polymerase pausing that is prominent in the GRO-seq protocol. However, the removal of the 200 bp sequence did not lead to a significant difference in results.) Reads were summarized to the rest of the gene body including intron regions in a strand-specific manner. Differential expression analysis was then performed using R [https://www.R-project.org/] and the limma+voom pipeline, consisting of TMM normalization using edgeR, statistical testing, and False Discovery Rate was controlled using the Benjamini-Hochberg procedure. Genes were filtered using fold changes and p values (fc ≥ 1.2 or fc ≥ 2 as described, and p ≤ 0.05 or including 0.05 ≤ p ≤ 0.07 as described). Both RNA-seq and GRO-seq analyses were performed by University of California Davis, Bioinformatics Core.

### Pathway analyses

We used the DAVID functional annotation tool^49^ as well as Enrichr for pathway analyses (KEGG, WikiPathway, Hallmark and Reactome)^158–160^.

### Fisher’s exact test

1. To determine if there is a statistically significant association between SATB1-upregulated genes with any of the TNBC subtypes (BL1, BL2, M, and LAR), we employed two-sided Fisher’s exact test. As universe of genes, the total set of genes being considered, represent all 1292 genes that were differentially expressed (SATB1-upregulated in the GRO-seq assay (FC1), with fc ≥ 1.2, and p ≤ 0.07. This is our “background” for enrichment analysis. SATB1-upregulated EMT genes (98 genes) were compared with TNBC-subtype-linked SATB1 upregulated genes (a), among which X number of genes (b) that overlap with the 98 genes using the 2x2 contingency table (shown below). We referred to Lehmann et al^115^ for all TNBC subtype-linked genes.

| All SATB1-dependent genes | EMT genes | Non-EMT genes | TOTAL |
| --- | --- | --- | --- |
| TNBC subtype-linked | a | b-a | b |
| Non-TNBC subtype-linked | 98-a=c | (1292-b)-c | 1292-b |
| Total | 98 | 1194 | 1292 |

- BL1-linked SATB1-upregulated genes (b=199 genes) among which 18 genes (a) overlap with the 98 EMT genes. The 2x2 table for BL1 is shown below.

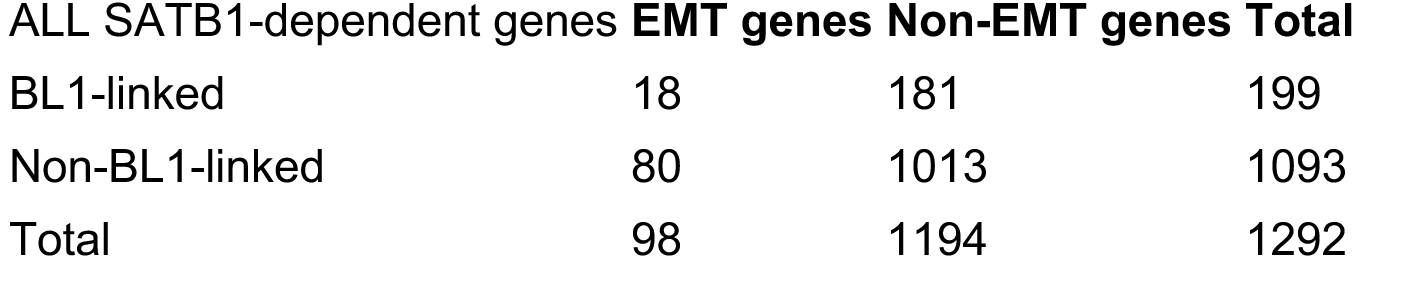

**Odds Ratio (OR)=1.25, Two-sided Fisher’s exact test p value=0.34**

- BL2-linked SATB1-upregulated genes, 296 genes(b), among which 48 genes (a) overlap with the 98 EMT genes.

**Odds Ratio (OR)=3.66, Two-sided Fisher’s exact test p value=4.65x10^-9^**

- M-linked SATB1-upregulated genes, 232 genes (b) among which 28 genes (a) overlap with the 98 EMT genes.

**Odds Ratio (OR)=1.94, Two-sided Fisher’s exact test p value=0.0035**

- LAR-linked SATB1-upregulated genes, 336 genes (b) among which 29 genes (a) overlap with the 98 EMT genes.

**Odds Ratio (OR)=1.21, Two-sided Fisher’s exact test p value=0.40**

2. To determine if **non-SATB1**-dependent EMT genes are significantly associated with any of the TNBC subtypes, the universe of genes in this case is all SATB1-independent genes among the 15101 genes minus SATB1-dependent gene (1292). 15101-1292=13809 genes. Among 300 EMT upregulated genes, 300-98=202 are non-SATB1-dependent EMT genes and the 202 genes are compared with TNBC subtype linked non-SATB1-upregulated genes (b), among which X number of overlap (a) with 202 EMT genes. The 2x2 contingent table used is shown below.

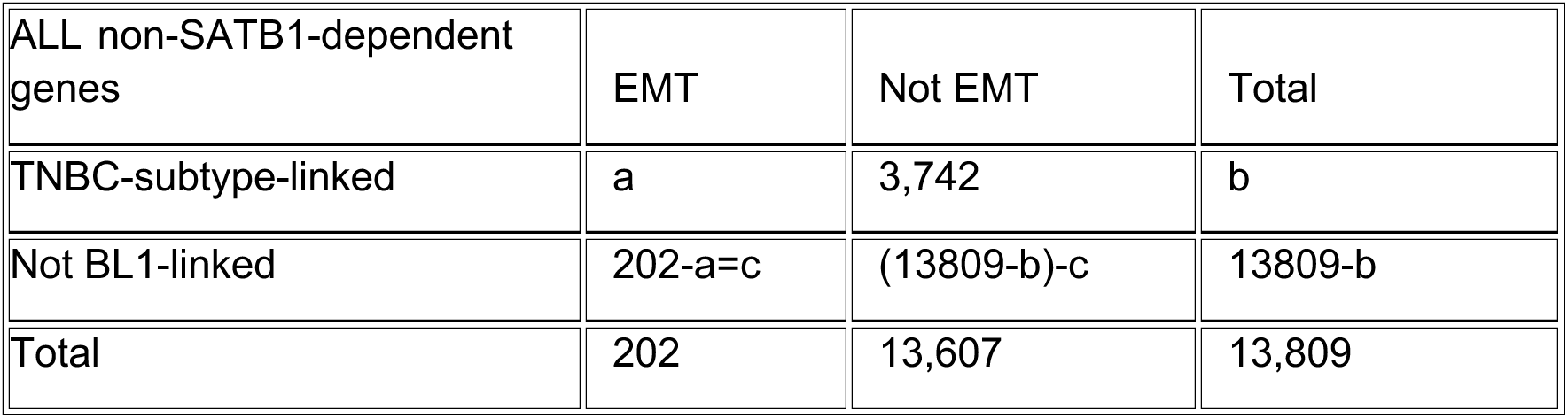

| ALL non-SATB1-dependent genes | EMT | Not EMT | Total |
| --- | --- | --- | --- |
| TNBC-subtype-linked | a | 3,742 | b |
| Not BL1-linked | $202-a=c$ | $(13809-b)-c$ | $13809-b$ |
| Total | 202 | 13,607 | 13,809 |

BL1-linked non-SATB1-dependent differentially expressed genes 3792 genes) among which 50 overlap with 202 EMT genes. The 2x2 table for BL1 subtype is shown below as an example.

| ALL non-SATB1-dependent genes | EMT | Not EMT | Total |
| --- | --- | --- | --- |
| BL1-linked | 50 | 3,742 | 3,792 |
| Not BL1-linked | 152 | 9,865 | 10,017 |
| Total | 202 | 13,607 | 13,809 |

**Odds ratio (OR)= 0.86, Two-sided Fisher’s exact p value=0.38**

- BL2-linked non-SATB1-dependent differentially expressed genes (3003 genes) among which 48 overlap with 202 EMT genes.

**Odds ratio (OR) = 1.09, Two-sided Fisher’s exact p-value = 0.61**

- M-linked non-SATB1-dependent differentially expressed genes (2967 genes) among which 72 overlap with 202 EMT genes.

**Odds ratio (OR) = 2.03, Two-sided Fisher’s exact p-value = 4.1 × 10⁻⁶**

- LAR-linked non-SATB1-dependent differentially expressed genes (4046 genes) among which 36 overlap with 202 EMT genes.

**Odds ratio (OR) = 0.51, Two-sided Fisher’s exact p-value = 1.7 × 10⁻⁴**

SATB1-independent EMT genes are significantly **under-represented** in LAR

3. Fisher’s exact test to analyze if there is statistically significant enrichment of SATB1-upregulated EMT genes among B[a]P-upregulated EMT genes. In this analysis, we used the following information. The universe: 300 EMT genes, SATB1-upregulated EMT genes:98, B[a]P-upregulated EMT genes:72, Overlap:36. Based on this information, a two-sided Fisher’s exact test showed Odds ratio: 2.68, and P-value: 4.99 × 10⁻⁴. This indicates a highly significant enrichment of carcinogen-regulated genes among SATB1-regulated EMT genes.

4. Fisher’s exact test to determine if EMT genes are enriched among SATB1-upregulated genes. We used the following information. The university: 15,101, SATB1-upregulated genes: 1,292, total EMT genes: 300, SATB1-dependent EMT: 98. 2x2 table is show below.

|  | EMT | Not EMT | Total |
| --- | --- | --- | --- |
| SATB1-dependent | 98 | 1,194 | 1,292 |
| Non-SATB1-dependent | 202 | 13,607 | 13,809 |
| Total | 300 | 14,801 | 15,101 |

The results showed Odds Ratio (OR)= 5.53 and two-sided Fisher’s exact p = 1 × 10⁻³⁰, indicating that EMT genes are significantly enriched among SATB1-upregulated genes relative to all other genes (including SATB1-downregulated and non-SATB1 dependent genes).

### Gene Set Enrichment Analysis

Gene set enrichment analysis (GSEA) was performed using the Broad Institute preranked GSEA methodology. For each analysis, genes were ranked according to log2 fold change (log2FC) derived from RNA-seq or GRO-seq differential expression analyses. Ranked gene lists were ordered from highest to lowest log2FC and used as input for preranked GSEA. Gene sets to be tested were defined from curated or experimentally derived lists. Preranked GSEA was carried out using a weighted enrichment statistic (weight p = 1) with 10,000 gene-set permutations. The enrichment score (ES) was calculated as the maximum deviation from zero of a running-sum statistic, and negative enrichment scores were retained when the absolute value of the minimum running-sum exceeded the maximum, in accordance with Broad Institute guidelines. Normalized enrichment scores (NES) were computed by normalizing observed ES values to the mean of the corresponding same-sign null distribution. Nominal p-values were estimated from the permutation-derived null distributions using a one-sided test in the direction of the observed ES. False discovery rates (FDR) were calculated following Broad Institute procedures by comparing the observed NES values to pooled null NES distributions within each ranked list. GSEA enrichment plots were generated in the Broad GSEA Desktop style, displaying the running ES curve and gene-set hit positions as thin vertical tick marks in a separate “Hits” panel.

## Data availability

All sequencing data for RNA-seq and GRO-seq in this study was uploaded to NCBI Gene Expression Omnibus (GEO) and is available under the accession GSE285433.

## Author Contributions

The project was conceived by T. K-S and Y.K. Y.K. modified the GRO-seq protocol and produced RNA-seq and GRO-seq data. T. K-S, Y.K and Y.V. contributed to data analyses. B.H and M.G. assisted Y.K for GRO-seq library preparation, sample preparation for GRO-seq and RNA-seq, and on data processing and organization. D.H. played a key role in advocating for funding from the California Breast Cancer Research Program (CBCRP).

## Acknowledgements

We thank Ms. Jie Li and Mr. Bradley Jenner at the Bioinformatics Core at the University of California, Davis, for their help with statistical and bioinformatic analyses for GRO-seq and RNA-seq data, and Dr. Reuben Thomas at the Bioinformatics Core at the Gladstone Institutes and Dr. Wenbo Li at University of Texas Health Science Center Houston for consultation on GRO-seq results. We thank Mr. Justus Williams from Washington College by performing initial DAVID analysis.

## Abbreviations

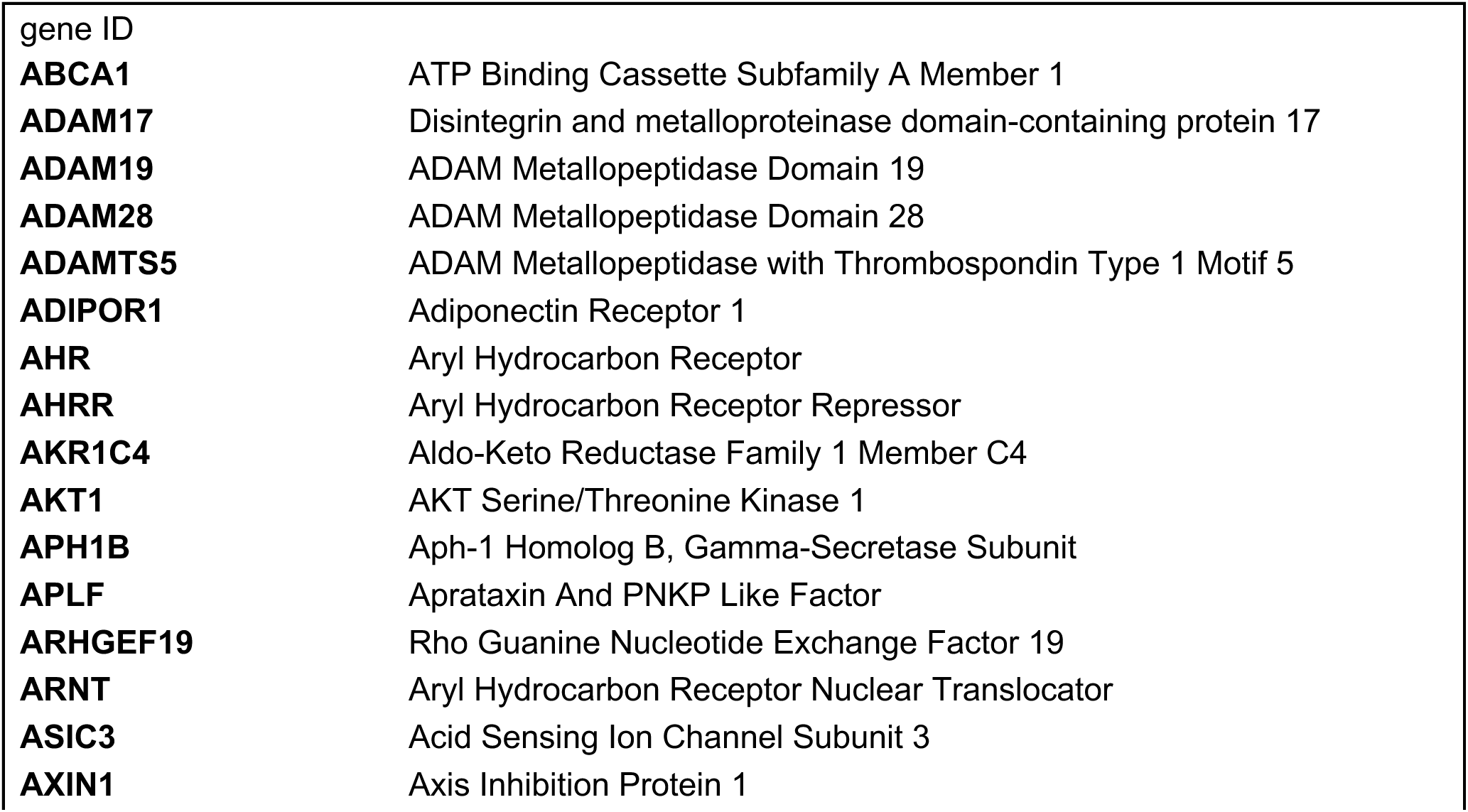
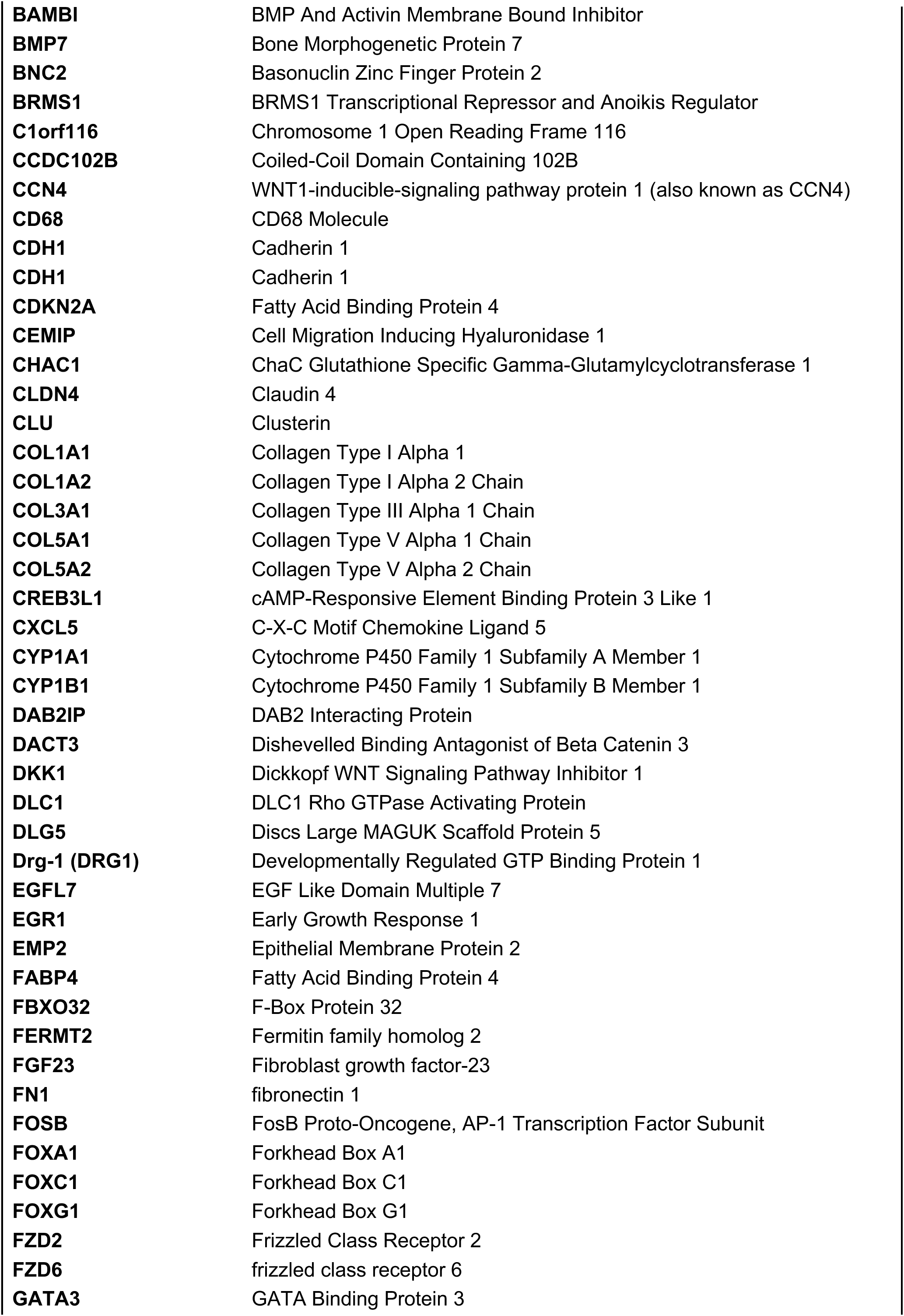
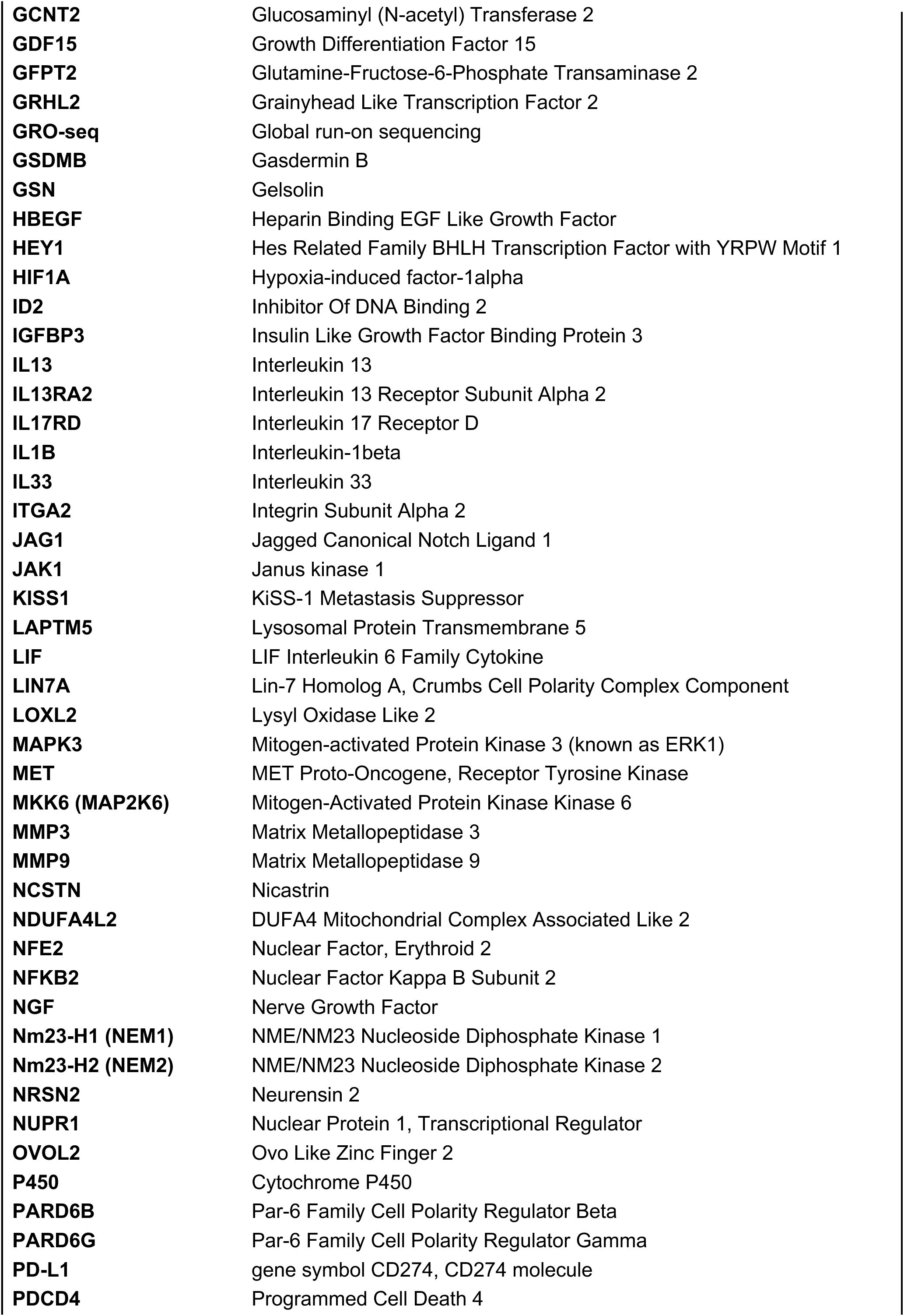
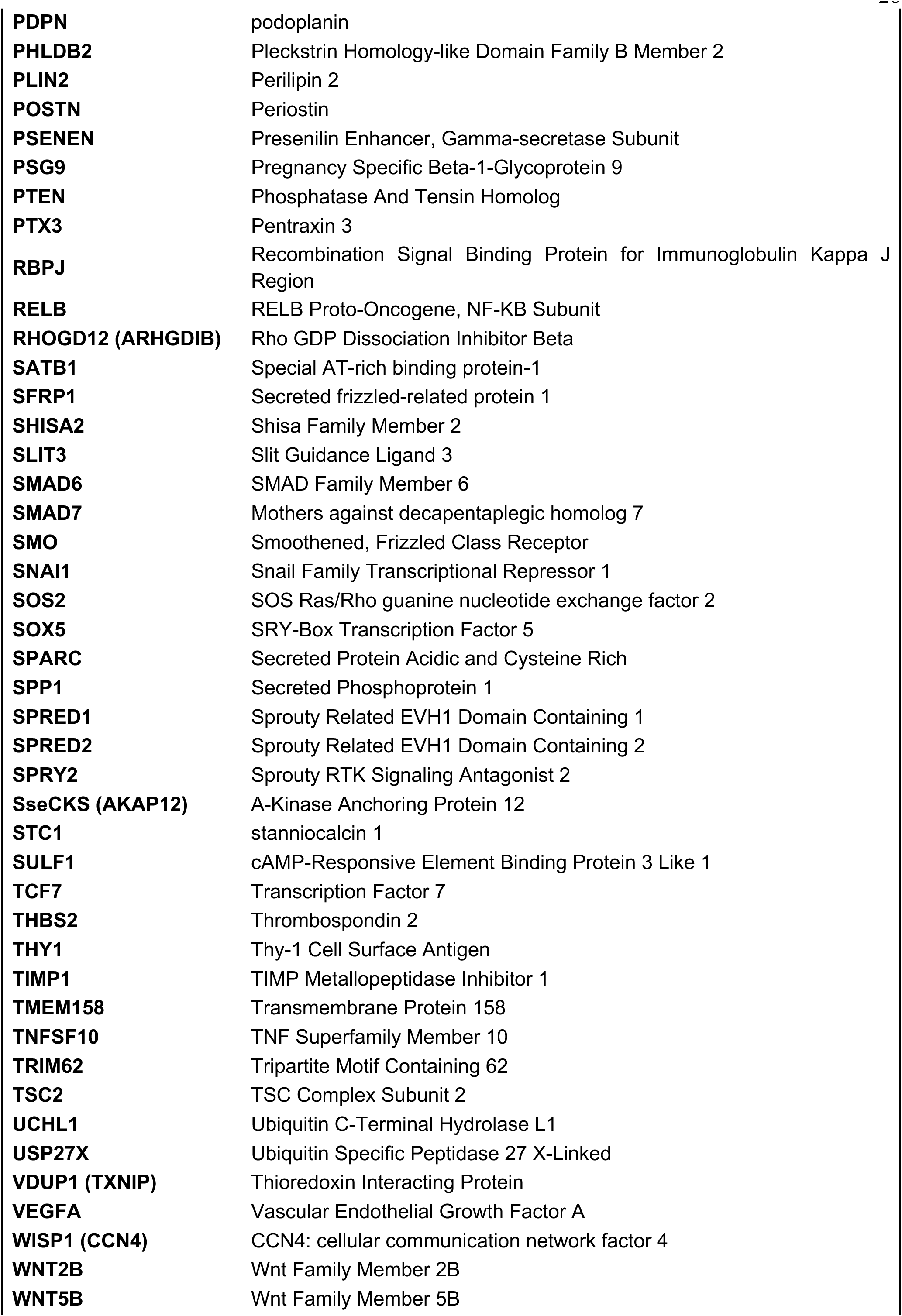
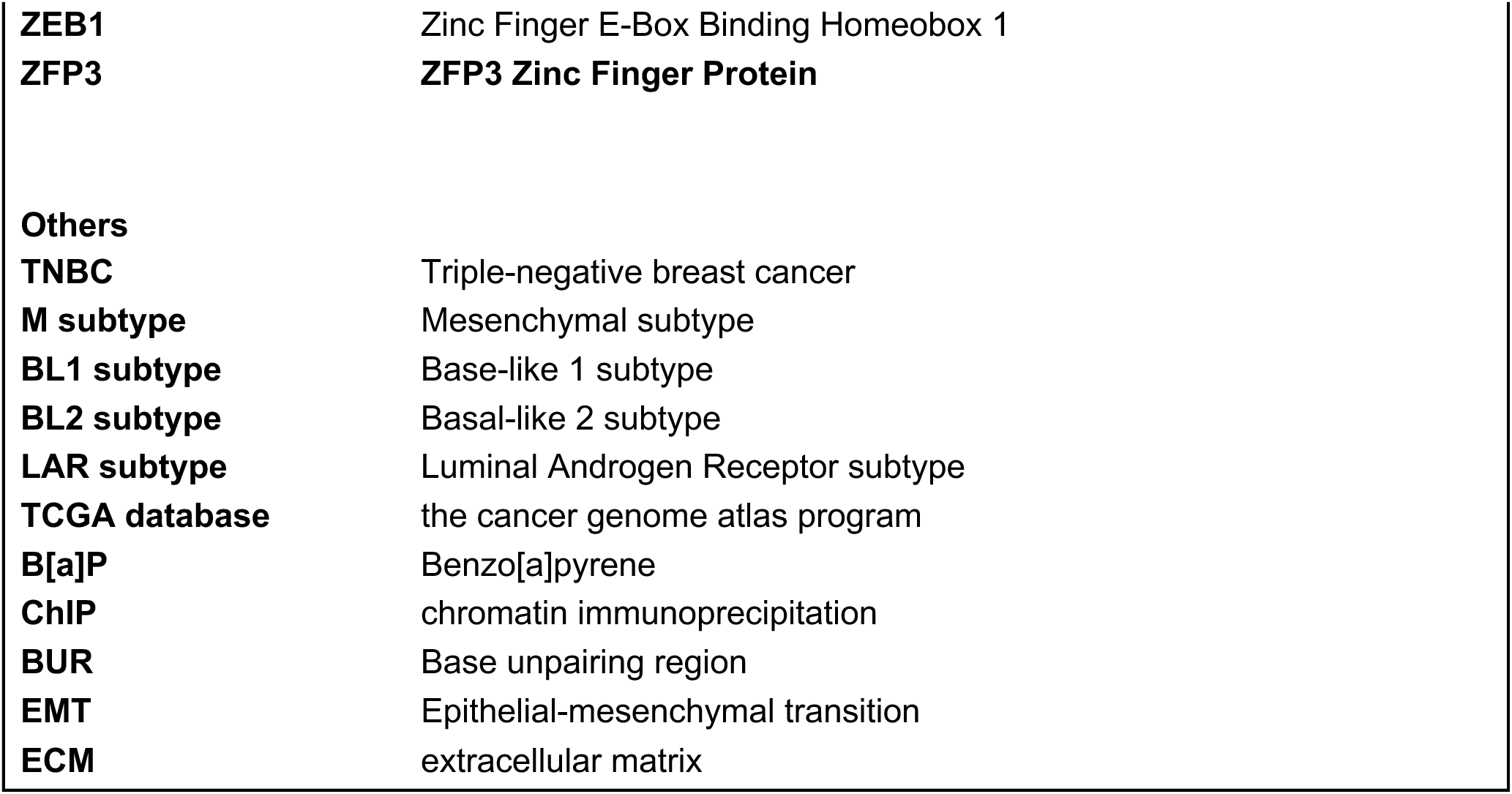

## References

1. Han, H.J., Russo, J., Kohwi, Y. & Kohwi-Shigematsu, T. SATB1 reprogrammes gene expression to promote breast tumour growth and metastasis. Nature 452, 187–193 (2008).

2. Ordinario, E., Han, H.J., Furuta, S., Heiser, L.M., Jakkula, L.R., Rodier, F., Spellman, P.T., Campisi, J., Gray, J.W., Bissell, M.J., Kohwi, Y. & Kohwi-Shigematsu, T. ATM suppresses SATB1-induced malignant progression in breast epithelial cells. PLoS One 7, e51786 (2012).

3. Kohwi-Shigematsu, T., Poterlowicz, K., Ordinario, E., Han, H.J., Botchkarev, V.A. & Kohwi, Y. Genome organizing function of SATB1 in tumor progression. Semin Cancer Biol 23, 72–79 (2013).

4. Li, Q.Q., Chen, Z.Q., Xu, J.D., Cao, X.X., Chen, Q., Liu, X.P. & Xu, Z.D. Overexpression and involvement of special AT-rich sequence binding protein 1 in multidrug resistance in human breast carcinoma cells. Cancer Sci 101, 80–86 (2010).

5. Liu, X., Zheng, Y., Qiao, C., Qv, F., Wang, J., Ding, B., Sun, Y. & Wang, Y. Expression of SATB1 and HER2 in breast cancer and the correlations with clinicopathologic characteristics. Diagn Pathol 10, 50 (2015).

6. Pan, Z., Jing, W., He, K., Zhang, L. & Long, X. SATB1 is Correlated with Progression and Metastasis of Breast Cancers: A Meta-Analysis. Cell Physiol Biochem 38, 1975–1983 (2016).

7. Fromberg, A., Engeland, K. & Aigner, A. The Special AT-rich Sequence Binding Protein 1 (SATB1) and its role in solid tumors. Cancer Lett 417, 96–111 (2018).

8. Glatzel-Plucinska, N., Piotrowska, A., Dziegiel, P. & Podhorska-Okolow, M. The Role of SATB1 in Tumour Progression and Metastasis. Int J Mol Sci 20 (2019).

9. Bai, J., Yang, G., Yu, Q., Chi, Q., Zeng, X. & Qi, W. SATB1 in cancer progression and metastasis: mechanisms and therapeutic potential. Front Oncol 15, 1535929 (2025).

10. Kohwi-Shigematsu, T. & Kohwi, Y. Torsional stress stabilizes extended base unpairing in suppressor sites flanking immunoglobulin heavy chain enhancer. Biochemistry 29, 9551–9560 (1990).

11. Cai, S., Han, H.J. & Kohwi-Shigematsu, T. Tissue-specific nuclear architecture and gene expression regulated by SATB1. Nat Genet 34, 42–51 (2003).

12. Kohwi, Y., Wong, X., Grange, M., Sexton, T., Richards H.W. , Kitagawa, Y. , Sakaguci, S., Liang, Y-C., Chuong, C-M., Botchkarev, V.A. , Taniuchi, I., Reddy, K. L., Kohwi-Shigematsu, T. Genome organization 1 by SATB1 binding to base-unpairing regions (BURs) provides a scaffold for SATB1-regulated gene expression. eLIFE **eLife** 14:**RP105915** (2025).

13. Alvarez, J.D., Yasui, D.H., Niida, H., Joh, T., Loh, D.Y. & Kohwi-Shigematsu, T. The MAR-binding protein SATB1 orchestrates temporal and spatial expression of multiple genes during T-cell development. Genes Dev 14, 521–535 (2000).

14. Cai, S., Lee, C.C. & Kohwi-Shigematsu, T. SATB1 packages densely looped, transcriptionally active chromatin for coordinated expression of cytokine genes. Nat Genet 38, 1278–1288 (2006).

15. Hao, B., Naik, A.K., Watanabe, A., Tanaka, H., Chen, L., Richards, H.W., Kondo, M., Taniuchi, I., Kohwi, Y., Kohwi-Shigematsu, T. & Krangel, M.S. An anti-silencer- and SATB1-dependent chromatin hub regulates Rag1 and Rag2 gene expression during thymocyte development. J Exp Med 212, 809–824 (2015).

16. Kitagawa, Y., Ohkura, N., Kidani, Y., Vandenbon, A., Hirota, K., Kawakami, R., Yasuda, K., Motooka, D., Nakamura, S., Kondo, M., Taniuchi, I., Kohwi-Shigematsu, T. & Sakaguchi, S. Guidance of regulatory T cell development by Satb1-dependent super-enhancer establishment. Nat Immunol 18, 173–183 (2017).

17. Kakugawa, K., Kojo, S., Tanaka, H., Seo, W., Endo, T.A., Kitagawa, Y., Muroi, S., Tenno, M., Yasmin, N., Kohwi, Y., Sakaguchi, S., Kowhi-Shigematsu, T. & Taniuchi, I. Essential Roles of SATB1 in Specifying T Lymphocyte Subsets. Cell Rep 19, 1176–1188 (2017).

18. Yasuda, K., Kitagawa, Y., Kawakami, R., Isaka, Y., Watanabe, H., Kondoh, G., Kohwi-Shigematsu, T., Sakaguchi, S. & Hirota, K. Satb1 regulates the effector program of encephalitogenic tissue Th17 cells in chronic inflammation. Nat Commun 10, 549 (2019).

19. Balamotis, M.A., Tamberg, N., Woo, Y.J., Li, J., Davy, B., Kohwi-Shigematsu, T. & Kohwi, Y. Satb1 ablation alters temporal expression of immediate early genes and reduces dendritic spine density during postnatal brain development. Mol Cell Biol 32, 333–347 (2012).

20. Close, J., Xu, H., De Marco Garcia, N., Batista-Brito, R., Rossignol, E., Rudy, B. & Fishell, G. Satb1 is an activity-modulated transcription factor required for the terminal differentiation and connectivity of medial ganglionic eminence-derived cortical interneurons. J Neurosci 32, 17690–17705 (2012).

21. Fessing, M.Y., Mardaryev, A.N., Gdula, M.R., Sharov, A.A., Sharova, T.Y., Rapisarda, V., Gordon, K.B., Smorodchenko, A.D., Poterlowicz, K., Ferone, G., Kohwi, Y., Missero, C., Kohwi-Shigematsu, T. & Botchkarev, V.A. p63 regulates Satb1 to control tissue-specific chromatin remodeling during development of the epidermis. J Cell Biol 194, 825–839 (2011).

22. Satoh, Y., Yokota, T., Sudo, T., Kondo, M., Lai, A., Kincade, P.W., Kouro, T., Iida, R., Kokame, K., Miyata, T., Habuchi, Y., Matsui, K., Tanaka, H., Matsumura, I., Oritani, K., Kohwi-Shigematsu, T. & Kanakura, Y. The Satb1 protein directs hematopoietic stem cell differentiation toward lymphoid lineages. Immunity 38, 1105–1115 (2013).

23. Skowronska-Krawczyk, D., Ma, Q., Schwartz, M., Scully, K., Li, W., Liu, Z., Taylor, H., Tollkuhn, J., Ohgi, K.A., Notani, D., Kohwi, Y., Kohwi-Shigematsu, T. & Rosenfeld, M.G. Required enhancer-matrin-3 network interactions for a homeodomain transcription program. Nature 514, 257–261 (2014).

24. Doi, Y., Yokota, T., Satoh, Y., Okuzaki, D., Tokunaga, M., Ishibashi, T., Sudo, T., Ueda, T., Shingai, Y., Ichii, M., Tanimura, A., Ezoe, S., Shibayama, H., Kohwi-Shigematsu, T., Takeda, J., Oritani, K. & Kanakura, Y. Variable SATB1 Levels Regulate Hematopoietic Stem Cell Heterogeneity with Distinct Lineage Fate. Cell Rep 23, 3223–3235 (2018).

25. Zhang, Y., Zheng, L., Le, M., Nakano, Y., Chan, B., Huang, Y., Torbaty, P.M., Kohwi, Y., Marcucio, R., Habelitz, S., Den Besten, P.K. & Kohwi-Shigematsu, T. SATB1 establishes ameloblast cell polarity and regulates directional amelogenin secretion for enamel formation. BMC Biol 17, 104 (2019).

26. Ribatti, D., Tamma, R. & Annese, T. Epithelial-Mesenchymal Transition in Cancer: A Historical Overview. Transl Oncol 13, 100773 (2020).

27. Thiery, J.P. Epithelial-mesenchymal transitions in tumour progression. Nat Rev Cancer 2, 442–454 (2002).

28. Tanabe, S., Quader, S., Cabral, H. & Ono, R. Interplay of EMT and CSC in Cancer and the Potential Therapeutic Strategies. Front Pharmacol 11, 904 (2020).

29. Terry, S., Savagner, P., Ortiz-Cuaran, S., Mahjoubi, L., Saintigny, P., Thiery, J.P. & Chouaib, S. New insights into the role of EMT in tumor immune escape. Mol Oncol 11, 824–846 (2017).

30. Huang, Y., Hong, W. & Wei, X. The molecular mechanisms and therapeutic strategies of EMT in tumor progression and metastasis. Journal of hematology & oncology 15, 129 (2022).

31. Lu, W. & Kang, Y. Epithelial-Mesenchymal Plasticity in Cancer Progression and Metastasis. Dev Cell 49, 361–374 (2019).

32. Yang, C., Tang, X., Guo, X., Niikura, Y., Kitagawa, K., Cui, K., Wong, S.T., Fu, L. & Xu, B. Aurora-B mediated ATM serine 1403 phosphorylation is required for mitotic ATM activation and the spindle checkpoint. Mol Cell 44, 597–608 (2011).

33. Bueno, R.C., Canevari, R.A., Villacis, R.A., Domingues, M.A., Caldeira, J.R., Rocha, R.M., Drigo, S.A. & Rogatto, S.R. ATM down-regulation is associated with poor prognosis in sporadic breast carcinomas. Ann Oncol 25, 69–75 (2014).

34. Feng, X., Li, H., Dean, M., Wilson, H.E., Kornaga, E., Enwere, E.K., Tang, P., Paterson, A., Lees-Miller, S.P., Magliocco, A.M. & Bebb, G. Low ATM protein expression in malignant tumor as well as cancer-associated stroma are independent prognostic factors in a retrospective study of early-stage hormone-negative breast cancer. Breast Cancer Res 17, 65 (2015).

35. Mastrangelo, G., Fadda, E. & Marzia, V. Polycyclic aromatic hydrocarbons and cancer in man. Environ Health Perspect 104, 1166–1170 (1996).

36. Phillips, D.H. Polycyclic aromatic hydrocarbons in the diet. Mutat Res 443, 139–147 (1999).

37. Li, X., Shen, C., Liu, X., He, J., Ding, Y., Gao, R., Mu, X., Geng, Y., Wang, Y. & Chen, X. Exposure to benzo[a]pyrene impairs decidualization and decidual angiogenesis in mice during early pregnancy. Environ Pollut 222, 523–531 (2017).

38. Archibong, A.E., Ramesh, A., Inyang, F., Niaz, M.S., Hood, D.B. & Kopsombut, P. Endocrine disruptive actions of inhaled benzo(a)pyrene on ovarian function and fetal survival in fisher F-344 adult rats. Reprod Toxicol 34, 635–643 (2012).

39. Perera, F.P., Estabrook, A., Hewer, A., Channing, K., Rundle, A., Mooney, L.A., Whyatt, R. & Phillips, D.H. Carcinogen-DNA adducts in human breast tissue. Cancer Epidemiol Biomarkers Prev 4, 233–238 (1995).

40. Rundle, A., Tang, D., Hibshoosh, H., Schnabel, F., Kelly, A., Levine, R., Zhou, J., Link, B. & Perera, F. Molecular epidemiologic studies of polycyclic aromatic hydrocarbon-DNA adducts and breast cancer. Environ Mol Mutagen 39, 201–207 (2002).

41. Balansky, R., Ganchev, G., Iltcheva, M., Steele, V.E., D’Agostini, F. & De Flora, S. Potent carcinogenicity of cigarette smoke in mice exposed early in life. Carcinogenesis 28, 2236–2243 (2007).

42. Amadou, A., Praud, D., Coudon, T., Deygas, F., Grassot, L., Faure, E., Couvidat, F., Caudeville, J., Bessagnet, B., Salizzoni, P., Gulliver, J., Leffondre, K., Severi, G., Mancini, F.R. & Fervers, B. Risk of breast cancer associated with long-term exposure to benzo[a]pyrene (BaP) air pollution: Evidence from the French E3N cohort study. Environ Int 149, 106399 (2021).

43. Castillo-Sanchez, R., Garcia-Hernandez, A., Torres-Alamilla, P., Cortes-Reynosa, P., Candanedo-Gonzales, F. & Salazar, E.P. Benzo[a]pyrene promotes an epithelial-to-mesenchymal transition process in MCF10A cells and mammary tumor growth and brain metastasis in female mice. Mol Carcinog 63, 1319–1333 (2024).

44. Guo, J., Xu, Y., Ji, W., Song, L., Dai, C. & Zhan, L. Effects of exposure to benzo[a]pyrene on metastasis of breast cancer are mediated through ROS-ERK-MMP9 axis signaling. Toxicol Lett 234, 201–210 (2015).

45. Malik, D.E., David, R.M. & Gooderham, N.J. Mechanistic evidence that benzo[a]pyrene promotes an inflammatory microenvironment that drives the metastatic potential of human mammary cells. Arch Toxicol 92, 3223–3239 (2018).

46. Castillo-Sanchez, R., Villegas-Comonfort, S., Galindo-Hernandez, O., Gomez, R. & Salazar, E.P. Benzo-[a]-pyrene induces FAK activation and cell migration in MDA-MB-231 breast cancer cells. Cell Biol Toxicol 29, 303–319 (2013).

47. Limsakul, P., Choochuen, P., Jungrungrueang, T. & Charupanit, K. Prognostic Markers in Tyrosine Kinases Specific to Basal-like 2 Subtype of Triple-Negative Breast Cancer. Int J Mol Sci 25 (2024).

48. Lopes, R., Agami, R. & Korkmaz, G. GRO-seq, A Tool for Identification of Transcripts Regulating Gene Expression. Methods Mol Biol 1543, 45–55 (2017).

49. Sherman, B.T., Hao, M., Qiu, J., Jiao, X., Baseler, M.W., Lane, H.C., Imamichi, T. & Chang, W. DAVID: a web server for functional enrichment analysis and functional annotation of gene lists (2021 update). Nucleic Acids Res 50, W216–W221 (2022).

50. Parvez, A., Choudhary, F., Mudgal, P., Khan, R., Qureshi, K.A., Farooqi, H. & Aspatwar, A. PD-1 and PD-L1: architects of immune symphony and immunotherapy breakthroughs in cancer treatment. Front Immunol 14, 1296341 (2023).

51. Wang, Z., You, P., Yang, Z., Xiao, H., Tang, X., Pan, Y., Li, X. & Gao, F. PD-1/PD-L1 immune checkpoint inhibitors in the treatment of unresectable locally advanced or metastatic triple negative breast cancer: a meta-analysis on their efficacy and safety. BMC Cancer 24, 1339 (2024).

52. Yang, Y., Yan, X., Bai, X., Yang, J. & Song, J. Programmed cell death-ligand 2: new insights in cancer. Front Immunol 15, 1359532 (2024).

53. Steeg, P.S. Metastasis suppressors alter the signal transduction of cancer cells. Nat Rev Cancer 3, 55–63 (2003).

54. Steeg, P.S., Horak, C.E. & Miller, K.D. Clinical-translational approaches to the Nm23-H1 metastasis suppressor. Clin Cancer Res 14, 5006–5012 (2008).

55. Salimian, N., Peymani, M., Ghaedi, K., Hashemi, M. & Rahimi, E. Collagen 1A1 (COL1A1) and Collagen11A1(COL11A1) as diagnostic biomarkers in Breast, colorectal and gastric cancers. Gene 892, 147867 (2024).

56. Li, X., Jin, Y. & Xue, J. Unveiling Collagen’s Role in Breast Cancer: Insights into Expression Patterns, Functions and Clinical Implications. Int J Gen Med 17, 1773–1787 (2024).

57. Paul, A.M., George, B., Saini, S., Pillai, M.R., Toi, M., Costa, L. & Kumar, R. Delineation of Pathogenomic Insights of Breast Cancer in Young Women. Cells 11 (2022).

58. Shi, Y., Zheng, C., Jin, Y., Bao, B., Wang, D., Hou, K., Feng, J., Tang, S., Qu, X., Liu, Y., Che, X. & Teng, Y. Reduced Expression of METTL3 Promotes Metastasis of Triple-Negative Breast Cancer by m6A Methylation-Mediated COL3A1 Up-Regulation. Front Oncol 10, 1126 (2020).

59. Yang, F., Lin, L., Li, X., Wen, R. & Zhang, X. Silencing of COL3A1 represses proliferation, migration, invasion, and immune escape of triple negative breast cancer cells via down-regulating PD-L1 expression. Cell biology international 46, 1959–1969 (2022).

60. Deng, J., Jiang, R., Meng, E. & Wu, H. CXCL5: A coachman to drive cancer progression. Front Oncol 12, 944494 (2022).

61. Romero-Moreno, R., Curtis, K.J., Coughlin, T.R., Miranda-Vergara, M.C., Dutta, S., Natarajan, A., Facchine, B.A., Jackson, K.M., Nystrom, L., Li, J., Kaliney, W., Niebur, G.L. & Littlepage, L.E. The CXCL5/CXCR2 axis is sufficient to promote breast cancer colonization during bone metastasis. Nat Commun 10, 4404 (2019).

62. Labedz, W., Przybyla, A., Zimna, A., Dabrowski, M. & Kubaszewski, L. The Role of Cytokines in the Metastasis of Solid Tumors to the Spine: Systematic Review. Int J Mol Sci 24 (2023).

63. Halpern, J.L., Kilbarger, A. & Lynch, C.C. Mesenchymal stem cells promote mammary cancer cell migration in vitro via the CXCR2 receptor. Cancer Lett 308, 91–99 (2011).

64. Winkler, J., Abisoye-Ogunniyan, A., Metcalf, K.J. & Werb, Z. Concepts of extracellular matrix remodelling in tumour progression and metastasis. Nat Commun 11, 5120 (2020).

65. Fang, M., Zou, M., Deng, H., Li, X., Zhu, Y., Shi, W. & Yang, L. THBS2 as a Key Regulator in Gastrointestinal Tumors: from Molecular Mechanisms to Clinical Applications. Curr Oncol Rep 28, 7 (2026).

66. Iwane, K. et al. Targeting fibroblast derived thrombospondin 2 disrupts an immune-exclusionary environment at the tumor front in colorectal cancer. Nat Commun 16, 11590 (2025).

67. Dorafshan, S., Razmi, M., Safaei, S., Gentilin, E., Madjd, Z. & Ghods, R. Periostin: biology and function in cancer. Cancer Cell Int 22, 315 (2022).

68. Suzuki, H., Kaneko, M.K. & Kato, Y. Roles of Podoplanin in Malignant Progression of Tumor. Cells 11 (2022).

69. Efthymiou, G., Saint, A., Ruff, M., Rekad, Z., Ciais, D. & Van Obberghen-Schilling, E. Shaping Up the Tumor Microenvironment With Cellular Fibronectin. Front Oncol 10, 641 (2020).

70. Lin, T.C., Yang, C.H., Cheng, L.H., Chang, W.T., Lin, Y.R. & Cheng, H.C. Fibronectin in Cancer: Friend or Foe. Cells 9 (2019).

71. Apte, R.S., Chen, D.S. & Ferrara, N. VEGF in Signaling and Disease: Beyond Discovery and Development. Cell 176, 1248–1264 (2019).

72. Mahaki, H., Nobari, S., Tanzadehpanah, H., Babaeizad, A., Kazemzadeh, G., Mehrabzadeh, M., Valipour, A., Yazdinezhad, N., Manoochehri, H., Yang, P. & Sheykhhasan, M. Targeting VEGF signaling for tumor microenvironment remodeling and metastasis inhibition: Therapeutic strategies and insights. Biomed Pharmacother 186, 118023 (2025).

73. Acerbi, I., Cassereau, L., Dean, I., Shi, Q., Au, A., Park, C., Chen, Y.Y., Liphardt, J., Hwang, E.S. & Weaver, V.M. Human breast cancer invasion and aggression correlates with ECM stiffening and immune cell infiltration. Integr Biol (Camb) 7, 1120–1134 (2015).

74. Arnold, S.A. & Brekken, R.A. SPARC: a matricellular regulator of tumorigenesis. J Cell Commun Signal 3, 255–273 (2009).

75. Nagaraju, G.P., Dontula, R., El-Rayes, B.F. & Lakka, S.S. Molecular mechanisms underlying the divergent roles of SPARC in human carcinogenesis. Carcinogenesis 35, 967–973 (2014).

76. Zhu, A., Yuan, P., Du, F., Hong, R., Ding, X., Shi, X., Fan, Y., Wang, J., Luo, Y., Ma, F., Zhang, P., Li, Q. & Xu, B. SPARC overexpression in primary tumors correlates with disease recurrence and overall survival in patients with triple negative breast cancer. Oncotarget 7, 76628–76634 (2016).

77. Augoff, K., Hryniewicz-Jankowska, A., Tabola, R. & Stach, K. MMP9: A Tough Target for Targeted Therapy for Cancer. Cancers (Basel) 14 (2022).

78. Lu, P., Takai, K., Weaver, V.M. & Werb, Z. Extracellular matrix degradation and remodeling in development and disease. Cold Spring Harb Perspect Biol 3 (2011).

79. Cox, T.R. & Erler, J.T. Remodeling and homeostasis of the extracellular matrix: implications for fibrotic diseases and cancer. Dis Model Mech 4, 165–178 (2011).

80. Domanegg, K., Sleeman, J.P. & Schmaus, A. CEMIP, a Promising Biomarker That Promotes the Progression and Metastasis of Colorectal and Other Types of Cancer. Cancers (Basel) 14 (2022).

81. Siddhartha, R. & Garg, M. Interplay Between Extracellular Matrix Remodeling and Angiogenesis in Tumor Ecosystem. Mol Cancer Ther 22, 291–305 (2023).

82. Wu, J., Wen, T., Marzio, A., Song, D., Chen, S., Yang, C., Zhao, F., Zhang, B., Zhao, G., Ferri, A., Cheng, H., Ma, J., Ren, H., Chen, Q.Y., Yang, Y. & Qin, S. FBXO32-mediated degradation of PTEN promotes lung adenocarcinoma progression. Cell Death Dis 15, 282 (2024).

83. Mondal, M., Conole, D., Nautiyal, J. & Tate, E.W. UCHL1 as a novel target in breast cancer: emerging insights from cell and chemical biology. Br J Cancer 126, 24–33 (2022).

84. Lee, J.E., Lim, Y.H. & Kim, J.H. UCH-L1 and UCH-L3 regulate the cancer stem cell-like properties through PI3 K/Akt signaling pathway in prostate cancer cells. Anim Cells Syst (Seoul) 25, 312–322 (2021).

85. Luo, Y., He, J., Yang, C., Orange, M., Ren, X., Blair, N., Tan, T., Yang, J.M. & Zhu, H. UCH-L1 promotes invasion of breast cancer cells through activating Akt signaling pathway. J Cell Biochem 119, 691–700 (2018).

86. Liu, Y., Song, Y., Ye, M., Hu, X., Wang, Z.P. & Zhu, X. The emerging role of WISP proteins in tumorigenesis and cancer therapy. J Transl Med 17, 28 (2019).

87. Saxena, N., Banerjee, S., Sengupta, K., Zoubine, M.N. & Banerjee, S.K. Differential expression of WISP-1 and WISP-2 genes in normal and transformed human breast cell lines. Mol Cell Biochem 228, 99–104 (2001).

88. Chiang, K.C., Yeh, C.N., Chung, L.C., Feng, T.H., Sun, C.C., Chen, M.F., Jan, Y.Y., Yeh, T.S., Chen, S.C. & Juang, H.H. WNT-1 inducible signaling pathway protein-1 enhances growth and tumorigenesis in human breast cancer. Sci Rep 5, 8686 (2015).

89. Xie, D., Nakachi, K., Wang, H., Elashoff, R. & Koeffler, H.P. Elevated levels of connective tissue growth factor, WISP-1, and CYR61 in primary breast cancers associated with more advanced features. Cancer Res 61, 8917–8923 (2001).

90. Xiu, M.X., Liu, Y.M. & Kuang, B.H. The oncogenic role of Jagged1/Notch signaling in cancer. Biomed Pharmacother 129, 110416 (2020).

91. Jeng, K.S., Sheen, I.S., Leu, C.M., Tseng, P.H. & Chang, C.F. The Role of Smoothened in Cancer. Int J Mol Sci 21 (2020).

92. Pelullo, M., Zema, S., Nardozza, F., Checquolo, S., Screpanti, I. & Bellavia, D. Wnt, Notch, and TGF-beta Pathways Impinge on Hedgehog Signaling Complexity: An Open Window on Cancer. Front Genet 10, 711 (2019).

93. Zhang, J., Tian, X.J. & Xing, J. Signal Transduction Pathways of EMT Induced by TGF-beta, SHH, and WNT and Their Crosstalks. J Clin Med 5 (2016).

94. Brenet, M., Martinez, S., Perez-Nunez, R., Perez, L.A., Contreras, P., Diaz, J., Avalos, A.M., Schneider, P., Quest, A.F.G. & Leyton, L. Thy-1 (CD90)-Induced Metastatic Cancer Cell Migration and Invasion Are beta3 Integrin-Dependent and Involve a Ca(2+)/P2X7 Receptor Signaling Axis. Front Cell Dev Biol 8, 592442 (2020).

95. Sung, V.Y.C., Knight, J.F., Johnson, R.M., Stern, Y.E., Saleh, S.M., Savage, P., Monast, A., Zuo, D., Duhamel, S. & Park, M. Co-dependency for MET and FGFR1 in basal triple-negative breast cancers. NPJ Breast Cancer 7, 36 (2021).

96. Yang, X., Liao, H.Y. & Zhang, H.H. Roles of MET in human cancer. Clin Chim Acta 525, 69–83 (2022).

97. Wu, Q., You, L., Nepovimova, E., Heger, Z., Wu, W., Kuca, K. & Adam, V. Hypoxia-inducible factors: master regulators of hypoxic tumor immune escape. Journal of hematology & oncology 15, 77 (2022).

98. Semenza, G.L. Hypoxia-inducible factors in physiology and medicine. Cell 148, 399–408 (2012).

99. Liu, Z., Sanders, A.J., Liang, G., Song, E., Jiang, W.G. & Gong, C. Hey Factors at the Crossroad of Tumorigenesis and Clinical Therapeutic Modulation of Hey for Anticancer Treatment. Mol Cancer Ther 16, 775–786 (2017).

100. Zhu, Y., Wang, W. & Wang, X. Roles of transcriptional factor 7 in production of inflammatory factors for lung diseases. J Transl Med 13, 273 (2015).

101. Huber, M.A., Azoitei, N., Baumann, B., Grunert, S., Sommer, A., Pehamberger, H., Kraut, N., Beug, H. & Wirth, T. NF-kappaB is essential for epithelial-mesenchymal transition and metastasis in a model of breast cancer progression. J Clin Invest 114, 569–581 (2004).

102. Mohamed, A.H., Abdelrahman, A.E., Fahmy, M.M., Waley, A.B., Saleh, M., Abdelwanis, A.H., Khalil, A.I. & Fouad, E.M. Prognostic Interplay of Caveolin1 and FOXC1 in Early Nonmetastatic Triple Negative Breast Cancer Undergoing Neoadjuvant Chemotherapy. Appl Immunohistochem Mol Morphol (2025).

103. Ozturk, O.B. & Arslan, R. FOXC1 expression profile in invasive breast carcinomas and its relationship with prognostic parameters. Pol J Pathol 76, 79–86 (2025).

104. Xie, Y., Wang, X., Wang, W., Pu, N. & Liu, L. Epithelial-mesenchymal transition orchestrates tumor microenvironment: current perceptions and challenges. J Transl Med 23, 386 (2025).

105. Fukumura, D., Kloepper, J., Amoozgar, Z., Duda, D.G. & Jain, R.K. Enhancing cancer immunotherapy using antiangiogenics: opportunities and challenges. Nat Rev Clin Oncol 15, 325–340 (2018).

106. Bergers, G., Brekken, R., McMahon, G., Vu, T.H., Itoh, T., Tamaki, K., Tanzawa, K., Thorpe, P., Itohara, S., Werb, Z. & Hanahan, D. Matrix metalloproteinase-9 triggers the angiogenic switch during carcinogenesis. Nat Cell Biol 2, 737–744 (2000).

107. Romero, Y., Wise, R. & Zolkiewska, A. Proteolytic processing of PD-L1 by ADAM proteases in breast cancer cells. Cancer Immunol Immunother 69, 43–55 (2020).

108. Waldhauer, I., Goehlsdorf, D., Gieseke, F., Weinschenk, T., Wittenbrink, M., Ludwig, A., Stevanovic, S., Rammensee, H.G. & Steinle, A. Tumor-associated MICA is shed by ADAM proteases. Cancer Res 68, 6368–6376 (2008).

109. Jones, K., Ballesteros, A., Mentink-Kane, M., Warren, J., Rattila, S., Malech, H., Kang, E. & Dveksler, G. PSG9 Stimulates Increase in FoxP3+ Regulatory T-Cells through the TGF-beta1 Pathway. PLoS One 11, e0158050 (2016).

110. Liu, Y.Y., Zhang, S., Yu, T.J., Zhang, F.L., Yang, F., Huang, Y.N., Ma, D., Liu, G.Y., Shao, Z.M. & Li, D.Q. Pregnancy-specific glycoprotein 9 acts as both a transcriptional target and a regulator of the canonical TGF-beta/Smad signaling to drive breast cancer progression. Clin Transl Med 10, e245 (2020).

111. Panda, V.K., Mishra, B., Nath, A.N., Butti, R., Yadav, A.S., Malhotra, D., Khanra, S., Mahapatra, S., Mishra, P., Swain, B., Majhi, S., Kumari, K., Radharani, N.N.V. & Kundu, G.C. Osteopontin: A Key Multifaceted Regulator in Tumor Progression and Immunomodulation. Biomedicines 12 (2024).

112. Jie, H., Ma, W. & Huang, C. Diagnosis, Prognosis, and Treatment of Triple-Negative Breast Cancer: A Review. Breast Cancer (Dove Med Press) 17, 265–274 (2025).

113. Lin, N.U., Claus, E., Sohl, J., Razzak, A.R., Arnaout, A. & Winer, E.P. Sites of distant recurrence and clinical outcomes in patients with metastatic triple-negative breast cancer: high incidence of central nervous system metastases. Cancer 113, 2638–2645 (2008).

114. Wang, D., Yang, Y., Rong, W., Fan, L., Yang, L., Chen, M., Yang, H. & He, Y. Natural history and prognostic nomogram of untreated triple negative breast cancer based on SEER database. Sci Rep 15, 23347 (2025).

115. Lehmann, B.D., Colaprico, A., Silva, T.C., Chen, J., An, H., Ban, Y., Huang, H., Wang, L., James, J.L., Balko, J.M., Gonzalez-Ericsson, P.I., Sanders, M.E., Zhang, B., Pietenpol, J.A. & Chen, X.S. Multi-omics analysis identifies therapeutic vulnerabilities in triple-negative breast cancer subtypes. Nat Commun 12, 6276 (2021).

116. Burstein, M.D., Tsimelzon, A., Poage, G.M., Covington, K.R., Contreras, A., Fuqua, S.A., Savage, M.I., Osborne, C.K., Hilsenbeck, S.G., Chang, J.C., Mills, G.B., Lau, C.C. & Brown, P.H. Comprehensive genomic analysis identifies novel subtypes and targets of triple-negative breast cancer. Clin Cancer Res 21, 1688–1698 (2015).

117. Masuda, H., Baggerly, K.A., Wang, Y., Zhang, Y., Gonzalez-Angulo, A.M., Meric-Bernstam, F., Valero, V., Lehmann, B.D., Pietenpol, J.A., Hortobagyi, G.N., Symmans, W.F. & Ueno, N.T. Differential response to neoadjuvant chemotherapy among 7 triple-negative breast cancer molecular subtypes. Clin Cancer Res 19, 5533–5540 (2013).

118. Opitz, C.A., Holfelder, P., Prentzell, M.T. & Trump, S. The complex biology of aryl hydrocarbon receptor activation in cancer and beyond. Biochem Pharmacol 216, 115798 (2023).

119. Guengerich, F.P. & Shimada, T. Oxidation of toxic and carcinogenic chemicals by human cytochrome P-450 enzymes. Chem Res Toxicol 4, 391–407 (1991).

120. Liu, J., Shi, Y., Wu, M., Zhang, F., Xu, M., He, Z. & Tang, M. JAG1 enhances angiogenesis in triple-negative breast cancer through promoting the secretion of exosomal lncRNA MALAT1. Genes Dis 10, 2167–2178 (2023).

121. Zhi, S., Chen, C., Huang, H., Zhang, Z., Zeng, F. & Zhang, S. Hypoxia-inducible factor in breast cancer: role and target for breast cancer treatment. Front Immunol 15, 1370800 (2024).

122. Pennacchietti, S., Michieli, P., Galluzzo, M., Mazzone, M., Giordano, S. & Comoglio, P.M. Hypoxia promotes invasive growth by transcriptional activation of the met protooncogene. Cancer Cell 3, 347–361 (2003).

123. Babina, I.S. & Turner, N.C. Advances and challenges in targeting FGFR signalling in cancer. Nat Rev Cancer 17, 318–332 (2017).

124. Manore, S.G., Doheny, D.L., Wong, G.L. & Lo, H.W. IL-6/JAK/STAT3 Signaling in Breast Cancer Metastasis: Biology and Treatment. Front Oncol 12, 866014 (2022).

125. Baghaie, L., Bunsick, D.A., Aucoin, E.B., Skapinker, E., Yaish, A.M., Li, Y., Harless, W.W. & Szewczuk, M.R. Pro-Inflammatory Cytokines Transactivate Glycosylated Cytokine Receptors on Cancer Cells to Induce Epithelial-Mesenchymal Transition to the Metastatic Phenotype. Cancers (Basel) 17 (2025).

126. Notani, D., Gottimukkala, K.P., Jayani, R.S., Limaye, A.S., Damle, M.V., Mehta, S., Purbey, P.K., Joseph, J. & Galande, S. Global regulator SATB1 recruits beta-catenin and regulates T(H)2 differentiation in Wnt-dependent manner. PLoS Biol 8, e1000296 (2010).

127. Yasui, D., Miyano, M., Cai, S., Varga-Weisz, P. & Kohwi-Shigematsu, T. SATB1 targets chromatin remodelling to regulate genes over long distances. Nature 419, 641–645. (2002).

128. Griffith, B.D. & Frankel, T.L. The Aryl Hydrocarbon Receptor: Impact on the Tumor Immune Microenvironment and Modulation as a Potential Therapy. Cancers (Basel) 16 (2024).

129. Hager, G.L., McNally, J.G. & Misteli, T. Transcription dynamics. Mol Cell 35, 741–753 (2009).

130. Li, B., Carey, M. & Workman, J.L. The role of chromatin during transcription. Cell 128, 707–719 (2007).

131. Nagel, M. & Taatjes, D.J. Regulation of RNA polymerase II transcription through re-initiation and bursting. Mol Cell 85, 1907–1919 (2025).

132. Chalancon, G., Ravarani, C.N., Balaji, S., Martinez-Arias, A., Aravind, L., Jothi, R. & Babu, M.M. Interplay between gene expression noise and regulatory network architecture. Trends Genet 28, 221–232 (2012).

133. Forero-Quintero, L.S., Raymond, W., Handa, T., Saxton, M.N., Morisaki, T., Kimura, H., Bertrand, E., Munsky, B. & Stasevich, T.J. Live-cell imaging reveals the spatiotemporal organization of endogenous RNA polymerase II phosphorylation at a single gene. Nat Commun 12, 3158 (2021).

134. Dar, R.D., Razooky, B.S., Singh, A., Trimeloni, T.V., McCollum, J.M., Cox, C.D., Simpson, M.L. & Weinberger, L.S. Transcriptional burst frequency and burst size are equally modulated across the human genome. Proc Natl Acad Sci U S A 109, 17454–17459 (2012).

135. Bintu, B., Mateo, L.J., Su, J.H., Sinnott-Armstrong, N.A., Parker, M., Kinrot, S., Yamaya, K., Boettiger, A.N. & Zhuang, X. Super-resolution chromatin tracing reveals domains and cooperative interactions in single cells. Science 362 (2018).

136. Finn, E.H., Pegoraro, G., Shachar, S. & Misteli, T. Comparative analysis of 2D and 3D distance measurements to study spatial genome organization. Methods 123, 47–55 (2017).

137. Gabriele, M., Brandao, H.B., Grosse-Holz, S., Jha, A., Dailey, G.M., Cattoglio, C., Hsieh, T.S., Mirny, L., Zechner, C. & Hansen, A.S. Dynamics of CTCF- and cohesin-mediated chromatin looping revealed by live-cell imaging. Science 376, 496–501 (2022).

138. Kaern, M., Elston, T.C., Blake, W.J. & Collins, J.J. Stochasticity in gene expression: from theories to phenotypes. Nat Rev Genet 6, 451–464 (2005).

139. Little, S.C., Tikhonov, M. & Gregor, T. Precise developmental gene expression arises from globally stochastic transcriptional activity. Cell 154, 789–800 (2013).

140. Hubalek, M., Czech, T. & Muller, H. Biological Subtypes of Triple-Negative Breast Cancer. Breast Care (Basel) 12, 8–14 (2017).

141. Guillen, K.P. et al. A human breast cancer-derived xenograft and organoid platform for drug discovery and precision oncology. Nat Cancer 3, 232–250 (2022).

142. van ’t Veer, L.J., Dai, H., van de Vijver, M.J., He, Y.D., Hart, A.A., Mao, M., Peterse, H.L., van der Kooy, K., Marton, M.J., Witteveen, A.T., Schreiber, G.J., Kerkhoven, R.M., Roberts, C., Linsley, P.S., Bernards, R. & Friend, S.H. Gene expression profiling predicts clinical outcome of breast cancer. Nature 415, 530–536 (2002).

143. Tian, S., Roepman, P., Van’t Veer, L.J., Bernards, R., de Snoo, F. & Glas, A.M. Biological functions of the genes in the mammaprint breast cancer profile reflect the hallmarks of cancer. Biomark Insights 5, 129–138 (2010).

144. Syed, Y.Y. Oncotype DX Breast Recurrence Score((R)): A Review of its Use in Early-Stage Breast Cancer. Mol Diagn Ther 24, 621–632 (2020).

145. Carlson, J.J. & Roth, J.A. The impact of the Oncotype Dx breast cancer assay in clinical practice: a systematic review and meta-analysis. Breast Cancer Res Treat 141, 13–22 (2013).

146. Vasaikar, S.V., Deshmukh, A.P., den Hollander, P., Addanki, S., Kuburich, N.A., Kudaravalli, S., Joseph, R., Chang, J.T., Soundararajan, R. & Mani, S.A. EMTome: a resource for pan-cancer analysis of epithelial-mesenchymal transition genes and signatures. Br J Cancer 124, 259–269 (2021).

147. Zhao, M., Liu, Y., Zheng, C. & Qu, H. dbEMT 2.0: An updated database for epithelial-mesenchymal transition genes with experimentally verified information and precalculated regulation information for cancer metastasis. J Genet Genomics 46, 595–597 (2019).

148. Wu, J., He, J., Zhang, J., Ji, H., Wang, N., Ma, S., Yan, X., Gao, X., Du, J., Liu, Z. & Hu, S. Identification of EMT-Related Genes and Prognostic Signature With Significant Implications on Biological Properties and Oncology Treatment of Lower Grade Gliomas. Front Cell Dev Biol 10, 887693 (2022).

149. Genies, C., Maitre, A., Lefebvre, E., Jullien, A., Chopard-Lallier, M. & Douki, T. The extreme variety of genotoxic response to benzo[a]pyrene in three different human cell lines from three different organs. PLoS One 8, e78356 (2013).

150. Dobin, A., Davis, C.A., Schlesinger, F., Drenkow, J., Zaleski, C., Jha, S., Batut, P., Chaisson, M. & Gingeras, T.R. STAR: ultrafast universal RNA-seq aligner. Bioinformatics 29, 15–21 (2013).

151. Robinson, M.D., McCarthy, D.J. & Smyth, G.K. edgeR: a Bioconductor package for differential expression analysis of digital gene expression data. Bioinformatics 26, 139–140 (2010).

152. Ritchie, M.E., Phipson, B., Wu, D., Hu, Y., Law, C.W., Shi, W. & Smyth, G.K. limma powers differential expression analyses for RNA-sequencing and microarray studies. Nucleic Acids Res 43, e47 (2015).

153. Law, C.W., Chen, Y., Shi, W. & Smyth, G.K. voom: Precision weights unlock linear model analysis tools for RNA-seq read counts. Genome Biol 15, R29 (2014).

154. Hochberg, Y. & Benjamini, Y. More powerful procedures for multiple significance testing. Stat Med 9, 811–818 (1990).

155. Li, H. & Durbin, R. Fast and accurate long-read alignment with Burrows-Wheeler transform. Bioinformatics 26, 589–595 (2010).

156. Liao, Y., Smyth, G.K. & Shi, W. featureCounts: an efficient general purpose program for assigning sequence reads to genomic features. Bioinformatics 30, 923–930 (2014).

157. Chae, M., Danko, C.G. & Kraus, W.L. groHMM: a computational tool for identifying unannotated and cell type-specific transcription units from global run-on sequencing data. BMC Bioinformatics 16, 222 (2015).

158. Xie, Z., Bailey, A., Kuleshov, M.V., Clarke, D.J.B., Evangelista, J.E., Jenkins, S.L., Lachmann, A., Wojciechowicz, M.L., Kropiwnicki, E., Jagodnik, K.M., Jeon, M. & Ma’ayan, A. Gene Set Knowledge Discovery with Enrichr. Curr Protoc 1, e90 (2021).

159. Kuleshov, M.V., Jones, M.R., Rouillard, A.D., Fernandez, N.F., Duan, Q., Wang, Z., Koplev, S., Jenkins, S.L., Jagodnik, K.M., Lachmann, A., McDermott, M.G., Monteiro, C.D., Gundersen, G.W. & Ma’ayan, A. Enrichr: a comprehensive gene set enrichment analysis web server 2016 update. Nucleic Acids Res 44, W90–97 (2016).

160. Chen, E.Y., Tan, C.M., Kou, Y., Duan, Q., Wang, Z., Meirelles, G.V., Clark, N.R. & Ma’ayan, A. Enrichr: interactive and collaborative HTML5 gene list enrichment analysis tool. BMC Bioinformatics 14, 128 (2013).

